# Genetic-substructure and complex demographic history of South African Bantu speakers

**DOI:** 10.1101/2020.08.11.243840

**Authors:** Dhriti Sengupta, Ananyo Choudhury, Cesar Fortes-Lima, Shaun Aron, Gavin Whitelaw, Koen Bostoen, Hilde Gunnink, Natalia Chousou-Polydouri, Peter Delius, Stephen Tollman, F Gomez-Olive Casas, Shane Norris, Felistas Mashinya, Marianne Alberts, Scott Hazelhurst, Carina M. Schlebusch, Michèle Ramsay, as members and collaborators of AWI-Gen and the H3Africa Consortium

**Author notes:** AWI-Gen and H3Africa members. Authors contributed equally to the work. Corresponding author: Michèle Ramsay.

## Abstract

South Eastern Bantu-speaking (SEB) groups constitute more than 80% of the population in South Africa. Despite clear linguistic and geographic diversity, the genetic differences between these groups have not been systematically investigated. Based on genome-wide data of over 5000 individuals, representing eight major SEB groups, we provide strong evidence for fine-scale population structure that broadly aligns with geographic distribution and is also congruent with linguistic phylogeny (separation of Nguni, Sotho-Tswana and Tsonga speakers). Although differential Khoe-San admixture plays a key role, the structure persists after Khoe-San ancestry-masking. The timing of admixture, levels of sex-biased gene flow and population size dynamics also highlight differences in the demographic histories of individual groups. The comparisons with five Iron Age farmer genomes further support genetic continuity over ∼400 years in certain regions of the country. Simulated trait genome-wide association studies further show that the observed population structure could have major implications for biomedical genomics research in South Africa.

## Introduction

The archaeological record and rock art evidence trace the presence of San-like hunter-gatherer culture in Southern Africa to at least 20 to 40 thousand years ago (kya)^1,2, 3^. Three sets of migration events have dramatically reshaped the genetic landscape of this geographic region in the last two millennia. The first of these was a relatively small scale migration of East African pastoralists, who introduced pastoralism to Southern Africa ∼2 kya^4–7^. This population was subsequently assimilated by local Southern African San hunter-gatherer groups, forming a new population that was ancestral to the Khoekhoe herder populations^8–12^. Today, Southern African Khoe and San (K-S) populations collectively refer to hunter-gatherer (San) and herder (Khoekhoe) communities. While K-S groups are distributed over a large geographic area today (spanning the Northern Cape Province of South Africa, large parts of Namibia, Botswana and Southern Angola), these groups are scattered, small and marginalised^13,14^.

The introduction of pastoralism in the region was closely followed by the arrival of the second set of migrants i.e. the Bantu-speaking (BS) agro-pastoralists. The archaeological record suggests that ancestors of the current-day BS populations undertook different waves of migration instead of a single large-scale movement^15–17^. The earliest communities spread along the East coast to reach the KwaZulu-Natal South coast by the mid-fifth century AD while the final major episode of settlement is estimated to be around AD 1350^18,19^. These archaeologically distinct groups gradually spread across present-day South Africa, interacting to various degrees with the K-S groups, eventually giving rise to South Africa’s diverse BS communities. The third major movement into Southern Africa was during the colonial era in the last four centuries when European colonists settled the area. During this period slave trade introduced additional intercontinental gene flow giving rise to complex genomic admixture patterns in current-day Southern African populations^20–23^.

South Africa has 11 official languages of which nine are Bantu languages belonging to this family’s South Eastern branch. Within these nine languages two large sub-clusters are traditionally distinguished: Nguni (including Zulu, Xhosa, Swazi, and Ndebele) and Sotho-Tswana (including Sotho, Tswana, and Pedi). Venda and Tsonga tend to be seen as independent linguistic entities. While the genetic diversity of K-S and mixed ancestry groups has been widely investigated^24^, the genetic diversity of the SEB-speaking (referred henceforth as SEB) groups has not been systematically investigated. One of the very early studies based on the Y-chromosome and a few autosomal markers, which included almost all the main SEB groups and covered most of the provinces from South Africa, indicated the possibility of genetic structure within the SEB populations^25^. However, many subsequent studies using genome-wide datasets did not investigate genetic differentiation or population structure within SEB groups, which consequently lead to its consideration as a group without clear internal sub-structure^21,26^. Moreover, studies including multiple SEB groups were often limited in terms of sample size or SEB group diversity^22,27,28^.

Based on an analysis of 5056 individuals (AWI-Gen study) genotyped on the Illumina H3A-genotyping array (∼2.3M SNPs), we report the first systematic study on the population structure within South African SEB groups. Although the eight SEB groups studied here have very specific geographic distributions of linguistic majority areas (LMAs) within the country, for our study they were sampled at three sites; Soweto (SWT) in Gauteng, Agincourt (AGT) in Mpumalanga, and Dikgale (DKG) in Limpopo province (**Table 1 and Fig. 1a**). To our knowledge, this is by far the largest genetic dataset from Southern Africa, and also the first comprehensive investigation into the demographic history of SEB groups.

**Table 1.** Distribution of the South Eastern Bantu-speaking (SEB) group by centre and ethnicity. The three columns for each centre shows for each SEB group: the total number of samples (All), the number of unrelated samples (PIHAT<0.18) (UR) and the ethno-linguistically concordant (EC) samples (self-reported ethno-linguistic identity of a participant is same as the ethno-linguistic identity of at least five of the six parents and grandparents). The column “All” corresponds to the dataset AWI-S1, “UR” corresponds to AWI-S2 and “EC” corresponds to AWI-S3 (**Supplementary Table 13**).

| SEB group | Agincourt centre |  |  | Dikgale centre |  |  | Soweto centre |  |  | Total |  |  |
| --- | --- | --- | --- | --- | --- | --- | --- | --- | --- | --- | --- | --- |
|  | All | UR | EC | All | UR | EC | All | UR | EC | All | UR | EC |
| Pedi | 36 | 33 | 0 | 1106 | 924 | 812 | 109 | 108 | 39 | 1251 | 1065 | 851 |
| Sotho | 97 | 79 | 0 | 9 | 9 | 5 | 285 | 278 | 41 | 391 | 366 | 46 |
| Swazi | 88 | 70 | 19 | 2 | 1 | 0 | 56 | 55 | 11 | 146 | 126 | 30 |
| Tsonga | 1941 | 1487 | 1369 | 52 | 47 | 22 | 117 | 110 | 47 | 2110 | 1644 | 1438 |
| Tswana | 1 | 1 | 1 | 14 | 13 | 5 | 234 | 228 | 67 | 249 | 242 | 73 |
| Venda | 5 | 5 | 2 | 23 | 21 | 6 | 47 | 47 | 16 | 75 | 73 | 24 |
| Xhosa | 3 | 3 | 2 | 6 | 6 | 4 | 169 | 168 | 57 | 178 | 177 | 63 |
| Zulu | 58 | 46 | 12 | 10 | 8 | 5 | 588 | 572 | 160 | 656 | 626 | 177 |
| **Total** | 2229 | 1724 | 1405 | 1222 | 1029 | 859 | 1605 | 1566 | 438 | 5056 | 4319 | 2702 |

**Fig. 1.**
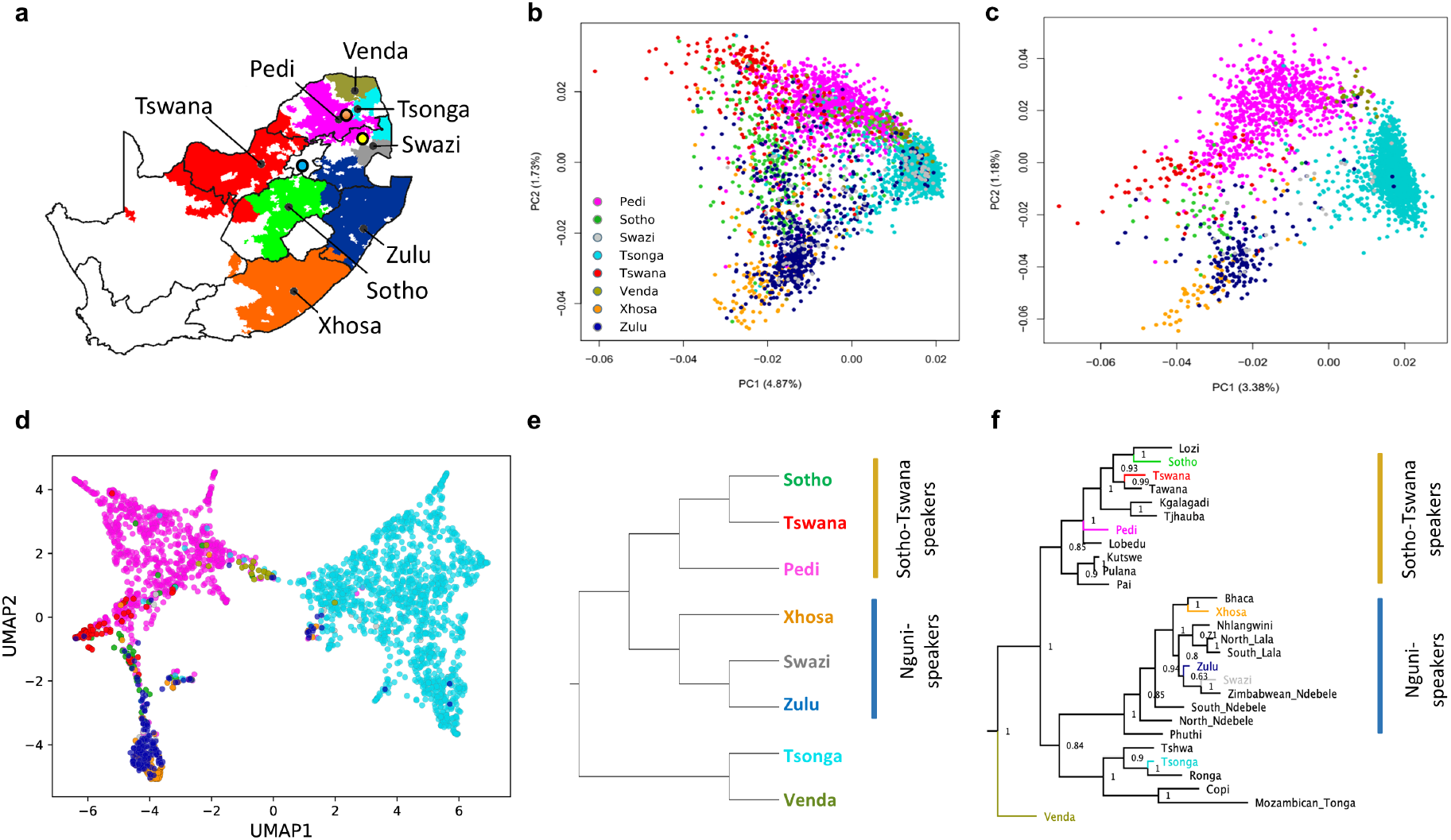
Population structure and genetic affinities of South Eastern Bantu-speaking (SEB) groups from South Africa correspond to both linguistic phylogeny and geographic distribution. **a**, Map showing the language majority areas (LMA) of each SEB group. The centroid of each of the regions is indicated using a black dot. The three sampling sites are shown in coloured circles; Soweto in blue, Dikgale in orange and Agincourt in yellow. **b**, Principal Component (PC) plot for the unrelated SEB samples (n=4,319) shows an overall correspondence between the distribution of SEB groups on the geographic map and the PCA. The colours showing the LMA for each SEB group on the geographic map corresponds to the colour used for the SEB group in the PCA. **c**, PC plot based on ethno-linguistically concordant samples (self-reported ancestry of the participant is the same as at least 5 of the parents and grandparents) (n= 2702) shows much clearer separation between the three major linguistic divisions (Sotho-Tswana, Nguni, and Xitsonga speakers). **d**, A composite representation of the first 10 PCs (generated using PCA-UMAP) also shows separation of the SEB-groups corresponding to the three major linguistic divisions. **e**, UPGMA tree based on pairwise F_ST_ distance between SEB groups. **f,** Linguistic phylogeny based on lexical data (majority-rule consensus tree with posterior probability values. The SEB groups from the current study are indicated using the same colours as used in the PCs. The topology of the trees in (**e)** and **(f)** shows an overall alignment.

Studies on population structure in South Africa should not be seen as justifying the ethnic nationalism generated by the country’s colonial and apartheid past. Our aim is to explore the role of genetic diversity in explaining population history and in health research. We recognise, and our study shows, that self-identity can involve considerable fluidity. We reject biological reductionist interpretations of our work.

## Results

### Fine-scale population structure within SEB

The principal component analysis (PCA) of unrelated SEB participants (AWI-S2 dataset) reflects the linguistic phylogeny with partial separation of Tsonga, Sotho-Tswana (Sotho, Pedi, Tswana) and Nguni (Zulu, Xhosa) speakers (**Fig. 1b**). The distribution of the SEB groups on the PC plot also largely mirrors the LMAs of these groups on the South African map (**Fig. 1a,b**) suggesting a correlation between genetic variation and geography.

The movement of populations from their LMAs to other regions during the last century is known to have enhanced the genetic exchange between different SEB groups, especially in urban areas such as Soweto^29^. These recent admixtures could result in incomplete boundaries observed between the SEB groups in the PC plot. In order to minimize the effect of such recent admixture on population structure detection, we performed PC analysis with 2,702 participants (**Fig. 1c**), who self-reported to share the same ethno-linguistic identity for at least five of the six parents and grandparents. We refer to these individuals as ethno-linguistically concordant (EC) participants hereafter (AWI-S3 dataset). The EC-based filtering step (**Table 1**) enhanced the resolution of these groups on the PC plot and also reduced the number of participants clustering with a different SEB group (**Fig. 1c**). PCA-UMAP analysis^30^, based on a composite of the first 10 PC coordinates estimated using EC participants, further illustrates the separation between the SEB groups (**Fig. 1d**). This highlights the importance of ethno-linguistically informed sampling for inferring the fine-scale population structure and also provides a possible rationale for why some previous studies, especially based on individuals from urban centres, could have underestimated population structure in SEB groups.

We compared our SEB populations to previously studied populations from Southern Africa^21,27,28,31^ by performing PC analysis with Merged dataset 2. The PC plot shows Zulu, Xhosa and Sotho individuals from these studies to group with corresponding SEB groups from the AWI-Gen study (**Supplementary Fig. 1**). Similarly, some of the individuals from Mozambique^31^ clustered close to Tsonga and Venda from our dataset, indicating the population structure to be largely robust. To avoid likely influence of sample size bias, we randomly downsized each group (AWI-S4 dataset), and the PC plots for this downsized data largely retained the fine-scale structure within SEB groups **(Supplementary Fig. 2)**.

Phylogenetic trees based on genetic distance (F_ST_) (**Fig. 1e, Supplementary Fig. 3a**) and linguistic phylogeny (**Fig. 1f, Supplementary Fig. 3b, Supplementary Note 1)** of the SEB groups shows overall alignment in topology. Similarly, the genetic (F_ST_) and geographical distances between the SEB groups also show a moderate correlation (Mantel test r value: 0.56, *P*-value=0.002). However, the overall low magnitude of F_ST_ values (**Supplementary Fig. 3c**) suggests that the fine-scale structure, although robust, corresponds to relatively small genetic distances.

### Differential K-S gene flow into various SEB groups

As K-S gene flow has been reported to be a major factor in differentiating SEB groups^22,27,28^, we estimated the level of K-S ancestry proportions in each SEB group (based on the Merged dataset 2 (EC downsized)) using an unsupervised clustering approach^32^. ADMIXTURE analysis at *K*=3 highlights the separation of African, K-S and Eurasian ancestry (blue, green and red component, respectively) **(Fig. 2a**). The various SEB groups showed differential levels of K-S gene flow varying from 1.5±2% in Tsonga to 20±6 % in Tswana (**Table 2**). The lowest cross-validation (CV) value was observed at *K*=5 which separates the Afro-Asiatic and the Central-West African ancestries (**Fig. 2a, Supplementary Fig. 4a**). Notably, the estimates show about 170 (4%) of the SEB participants harbour more than 5% Eurasian-like ancestry (**Table 2**). As there has been no systematic study to estimate the level of Eurasian ancestry in the more Northern provinces of the country, we were unable to estimate whether the observed level of Eurasian ancestry is common in SEB groups from these geographic areas or a cohort-specific feature. Our results could provide a baseline for future studies on Eurasian admixture in SEB groups.

**Fig. 2.**
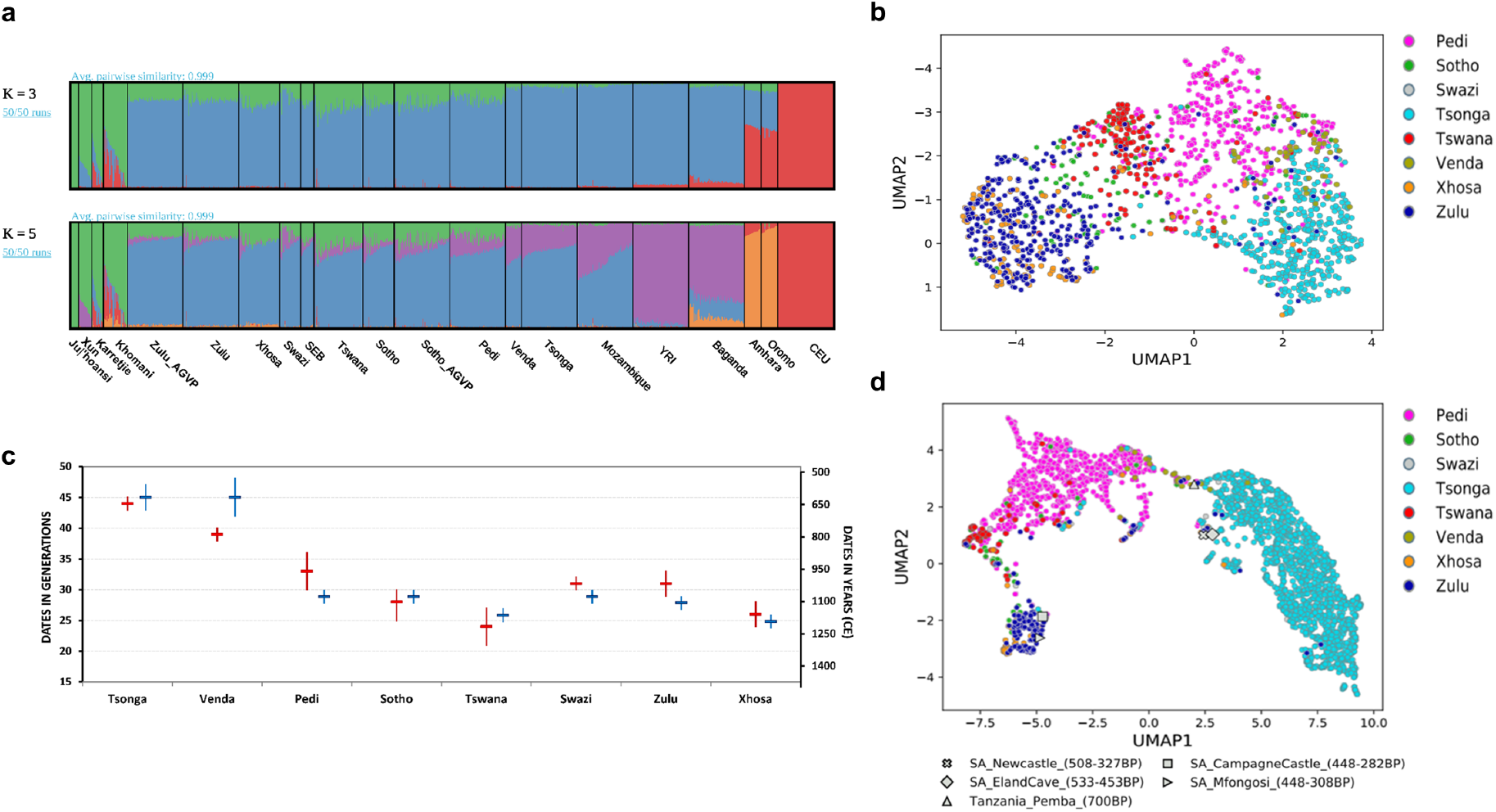
Gene flow into and genetic continuity of South Eastern Bantu-speaking (SEB) groups. **a**, ADMIXTURE plots (from *K*=3 to *K*=5) based on the merged dataset with downsized ethno-linguistically concordant individuals. At *K*=3, the plot shows differences in the level of Khoe-San (K-S) gene flow (shown in green) into different SEB groups, with Tswana and Xhosa showing the highest K-S ancestry proportion and Tsonga and Venda the lowest. Baganda (from Uganda); Amhara, Oromo and Somali (from Ethiopia); Sotho_AGVP and Zulu_AGVP (from South Africa) are from (ref. ^28^) datasets. The Yoruba (YRI) and Central European (CEU) are from the 1000 Genomes Project dataset^55^. **b**, Composite representation of the first 10 PCs (generated using PCA-UMAP) showing population structure in SEB groups persists even after K-S ancestry masking. **c**, Dates for K-S admixture in SEB populations estimated using fastGLOBETROTTER (red) and MALDER (blue). Figure also showing 95% CI bars (vertical lines) from each method. First y-axis shows admixture dates in generations ago, while in the second y-axis shows the actual estimated dates. CE refers to the Common Era. **d**, Composite representation of the first 10 PCs comparing Iron-Age genomes to our SEB groups indicate genetic continuity for the last few centuries in certain regions of South Africa.

**Table 2.** Ancestry proportions for various South Eastern Bantu-speaking (SEB) groups estimated using ADMIXTURE analysis (at *K*=3).

| SEB group | Bantu-related ancestry (%) |  | Khoen-San-related ancestry (%) |  | Eurasian-related ancestry (%) |  | Sample size |
| --- | --- | --- | --- | --- | --- | --- | --- |
|  | Mean | ±SD | Mean | ±SD | Mean | ±SD |  |
| Pedi | 88.28 | 5.11 | 10.61 | 4.71 | 1.12 | 2.83 | 1065 |
| Sotho | 84.17 | 8.36 | 14.65 | 7.40 | 1.18 | 3.60 | 366 |
| Tswana | 78.19 | 7.44 | 20.49 | 6.02 | 1.32 | 4.59 | 242 |
| Swazi | 90.43 | 8.02 | 8.69 | 7.49 | 0.87 | 2.59 | 126 |
| Xhosa | 80.24 | 5.90 | 17.62 | 4.86 | 2.14 | 3.17 | 177 |
| Zulu | 84.64 | 6.32 | 13.58 | 4.72 | 1.78 | 4.16 | 626 |
| Tsonga | 97.80 | 2.86 | 1.56 | 2.43 | 0.65 | 1.21 | 1644 |
| Venda | 91.31 | 6.99 | 6.45 | 5.15 | 2.24 | 4.16 | 73 |

ADMIXTURE analysis on the full set of unrelated samples (Merged dataset 1) detected considerable within-population variation in ancestry proportions for some of the SEB groups (**Supplementary Fig. 4b**). When partitioned by the study site, four of the SEB groups (Zulu, Sotho, Pedi and Swazi) show significantly higher K-S ancestry in participants originating from SWT in comparison to participants from AGT (**Supplementary Table 1, Supplementary Fig. 4c-f**). These differences for populations such as Swazi and Zulu were also distinguishable in a PC plot that includes the site of collection information along with group labels (**Supplementary Fig. 5**). These observations emphasize the importance of careful consideration of sampling locations in addition to ethno-linguistic concordance, for a comprehensive estimation of the fine-scale population structure.

We further investigated whether differential K-S gene flow was the only factor leading to the observed population structure, by masking non-Bantu-related haplotypes in each SEB individual (**see Methods**). Ancestry-specific PCA after masking haploid genomes shows that the core differences within SEB groups, although reduced, persist even after accounting for differential K-S gene flow (**Fig. 2b**, **Supplementary Fig. 6**). The observed structure between SEB groups could therefore be attributed to additional historical and demographic factors, such as multiple expansion movements into Southern Africa, different points of origin and isolation due to geography.

### Dating admixture events in SEB groups

To reconstruct the timeline of migration of each SEB group, we dated the admixture between the best BS and K-S source populations for each group using fastGLOBETROTTER^33^ (**Fig. 2c, Supplementary Table 2**). As the range of K-S populations is estimated to have been much wider in the past compared to their present distribution, some of these admixture events might have occurred beyond the boundaries of the country. Moreover, it is also possible that in some cases gene flow from the K-S might not have immediately followed the arrival of the BS populations. Nevertheless, it is reasonable to expect that major differences in admixture dating could be broadly indicative of the differences in dates of arrival and settlement of the ancestral SEB group in different regions of the country.

Consistent with many previous studies^22,28,34,35^, the inferred dating pattern indicates that the contact between the ancestors of all the SEB groups and K-S populations included in our study, occurred within the last 45 generations (∼1300 years). Moreover, for all SEB groups, a single admixture event model was detected to be the best-guess conclusion by fastGLOBETROTTER (**Supplementary Note 2)**. Tsonga and Venda show the oldest admixture dates (around 45 generations ago) while the admixture dates for the other SEB groups range between 24-33 generations ago. The presence of SEB groups on the South African landscape is assumed to date back to the fourth century AD, from which time there is considerable archaeological evidence for interaction with K-S that probably included admixture^36^. The admixture dates for Tsonga and Venda, therefore, suggest that these SEB groups of southern Mozambique and North-Eastern South Africa could be descendants from one of the earlier episodes of settlement in this region.

The admixture dates correlate broadly with geography, with more Northern populations showing relatively older dates compared to Southern populations, for example, Zulu compared to Xhosa **(Fig. 2c, Supplementary Table 2).** Even among the groups from the inland plateau region (referred to as the highveld), we observed more recent dates for more Southern/Western populations (the Sotho and Tswana), compared to the more Northern Pedi. However, we also observed exceptions to these trends, such as a large difference between the K-S admixture dates in geographically neighbouring Pedi and Tsonga. Multiple westward movements of Tsonga-speakers from Mozambique in the last few centuries have been reported^37^ suggesting that the Tsonga and Pedi might have been separated by much greater geographic distances in the past, likely explaining the stark differences in admixture dating.

To test the robustness of the observed dating patterns, we also dated these admixture events using MALDER^38^ (**Fig. 2c**) and MOSAIC^39^ (**Supplementary Table 2, Supplementary Note 2**). Although there are some differences in the predicted time-scales of admixture events obtained using these dating methods (MOSAIC for most groups generated younger dates), all the admixture dating methods demonstrate the same pattern (**Supplementary Table 2, Supplementary Fig. 7, Supplementary Note 2**). The estimated dates of Eurasian admixture in SEB groups (4-5 generations ago, **Supplementary Table 3**) is consistent with the rather recent settlement of European ancestry populations in the geographic region corresponding to the three sampling sites^40^.

### Relationship between ancient genomes and modern SEB groups

The availability of Iron Age genomes from Southern Africa provided us with the unique opportunity to compare the affinities of present-day SEB groups to populations living in these areas centuries ago^10,11^. The PCA and PCA-UMAP projecting five Iron Age Bantu-related genomes (300 to 700 years old) onto the genetic variation of present-day SEB individuals (**Fig. 2d, Supplementary Fig. 8**) show these genomes to be on a temporal cline with the older genomes (Pemba, Eland Cave and Newcastle; ranging ∼700-450 BP) appearing closer to the Tsonga and Venda, while more recent genomes (Champagne Castle and Mfongosi; ranging from ∼448-300 BP) occurring closer to the Nguni-speakers. This cline of the Iron Age genomes also aligns with geographic distribution from North to South, as well as increasing levels of K-S ancestry in them^10^. More ancient genomes from Southern Africa would be required to test whether the trends observed in these Iron Age genomes are indicative of phases in the movement of groups further South with time, a process marked by concomitant increase of K-S ancestry in the migrants. Interestingly, the wider geographic region of Northern KwaZulu Natal around Champagne Castle in Central-East South Africa, where the youngest of these Iron-Age genomes was collected, is still dominated by Nguni-speakers (**Fig. 1a**), providing support for at least four centuries of genetic continuity in certain regions of South Africa.

### Sex-specific admixture patterns

In accordance with several previous reports^27,41,42^, the comparison of mitochondrial DNA and Y chromosome haplogroup distributions (**Fig. 3a, Supplementary Table 4, Supplementary Table 5**) shows evidence for relatively higher maternal gene-flow from K-S into all the SEB groups (**Supplementary Note 3).** Comparison of the autosomal and X-chromosome contributions also supports K-S biased maternal gene-flow (Fig. 3b). However, the level of this bias varies widely between groups (**Fig. 3a,b, Supplementary Table 6**). The lack of any correlation between the extent of this bias and level of admixture/admixture dates suggests that the nature of interaction between K-S and BS could have been determined by other demographic factors (**Supplementary Note 3)**.

**Fig. 3.**
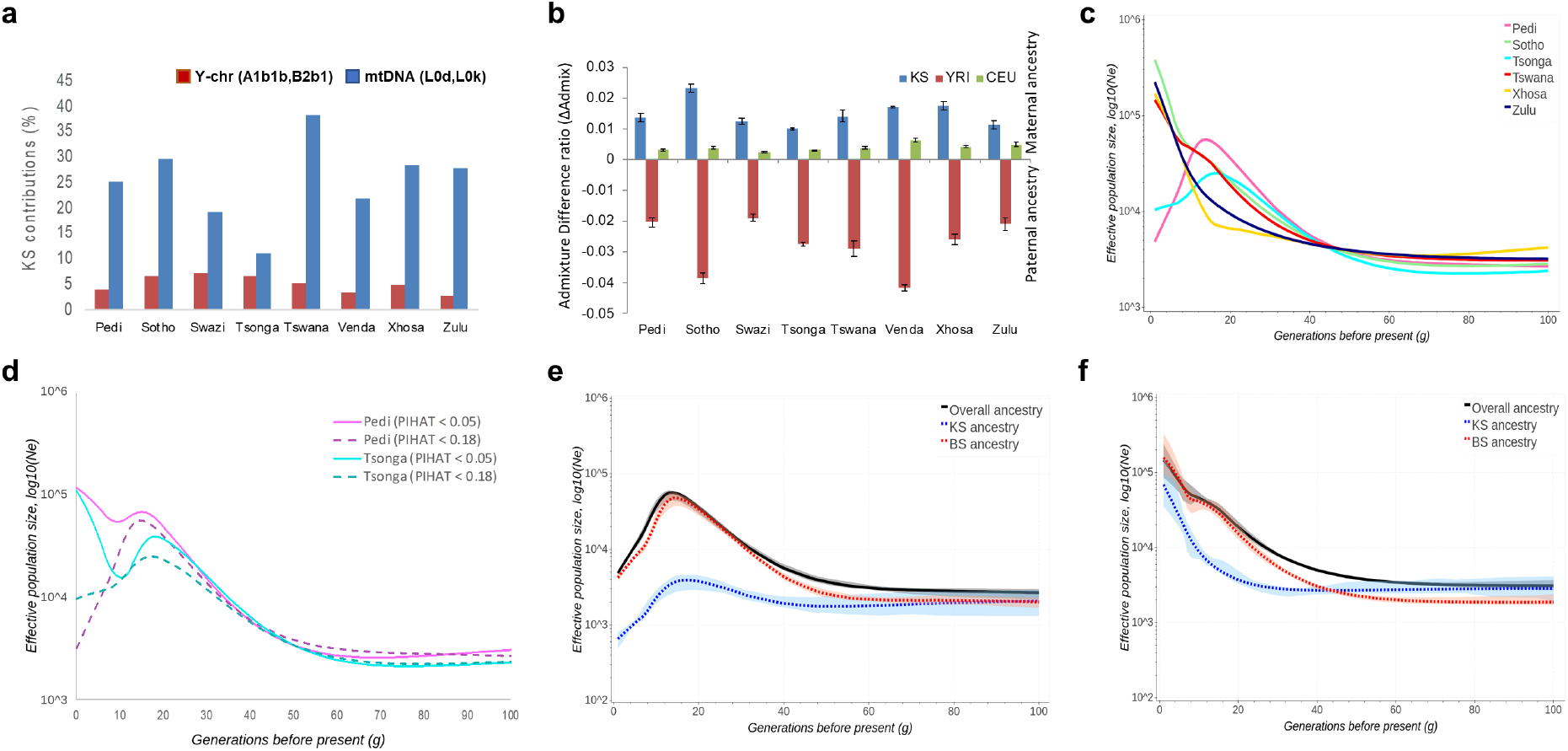
Insights into the demographic history of South Eastern Bantu-speaking (SEB) groups. **a**, Distribution of Khoe-San (K-S) associated mitochondrial and Y chromosome haplogroups in the SEB groups shows higher maternal contribution from K-S **b**, The analysis of admixture difference ratio (based on X chromosomal and autosomal contributions) confirms this trend and shows the level of bias to vary strongly between the SEB groups. The bars show admixture differences for the three contributing ancestries. Blue shows K-S, red shows Bantu-speaker (represented by KGP Yuroba (YRI)) and green shows Eurasian (represented by KGP Central European (CEU)) ancestries for each SEB group. Positive bar values denotes a maternal bias whereas negative values denotes paternal bias in contributions from an ancestry. **c**, Effective population size (*Ne*) fluctuations (estimated using IBDNe) shows SEB groups to differentiate mainly in the last 40 generations. **d**, *Ne* profile differences in Pedi and Tsonga before and after removal of individuals with 0.05<PIHAT<0.18. **e-f,** Ancestry-specific IBDNe (AS-IBDNe) based evaluation of the relative contribution of K-S and BS to the *Ne* profiles in **(e)** Pedi, and **(f)** Tswana. For **(e)** and **(f)**, the black line shows the overall (“true”) *Ne* while the red and blue lines shows the *Ne* for BS and K-S ancestral components, respectively. The shaded areas around each line demarcates 95% confidence intervals. The plots show the level of K-S ancestry to correlate with the extent of influence on overall *Ne*.

### Variation of effective population size through time

We investigated changes in the effective population size (*Ne*) of each SEB group over the last 100 generations by analysing the sharing patterns of identity-by-descent (IBD) segments using IBDNe^43^. As depicted in **Fig. 3c**, the *Ne* for all the SEB groups was very similar for the 100^th^ to the 40^th^ generations before present. It needs to be noted that most of the present-day SEB groups did not exist, as such, more than 50 generations ago and the older estimates here correspond to possible ancestral populations of these groups. The period of around 40 generations ago also corresponds to the estimated time scale for the oldest K-S admixture dates (**Fig. 2c**). From the 40^th^ generation onwards, the Nguni-speakers and Sotho-Tswana speakers start showing distinct and characteristic *Ne* profiles, which possibly reflect migration events that separated these populations in terms of geography. Similarly, the dates for the initiation of population size increase of the Zulu around 25 generations ago, broadly corresponds to the time (around AD 1300) when Nguni-speakers first began to move North-west into the interior, becoming the first BS in South Africa to occupy grasslands^44,45^. The comparison of Sotho and Zulu *Ne* profiles between our study and samples from a previous study^28^ shows a high concordance, demonstrating an overall robustness in these estimates (**Supplementary Fig. 9**).

A high level of cryptic relatedness (CR) in a population could strongly impact estimates based on IBD-sharing. Despite adopting a sampling strategy aimed at minimizing the recruitment of genetically related participants, we observed very high levels of CR in Tsonga and Pedi (**Figure 3d**, **Supplementary Fig. 10, Supplementary Note 4**). Notably, in contrast to other SEB groups, both Pedi and Tsonga showed a strong *Ne* decline in the last 20 generations which could be a function of CR. Therefore, we re-estimated the *Ne* profiles for these groups based only on unrelated participants with PIHAT<0.05 (**Fig. 3d**). The filtering for relatedness removed the recent drop in population size observed in both populations. The *Ne* profile for Pedi participants after filtering also shows much higher resemblance to other Sotho-Tswana speakers. However, whether the related or the unrelated samples represent the actual demographic history of these SEB groups remains an open question for future studies.

We further partitioned the contribution of the two major source ancestries (K-S and BS) to the *Ne* profiles of the SEB groups by using the ancestry-specific (AS) IBD-Ne approach^43^. The results depicted in **Fig. 3e, 3f and Supplementary Fig. 11** clearly show that the *Ne* curves, although being driven by BS ancestry, are also affected by K-S gene flow. The K-S ancestry impact on the *Ne* profiles was correlated with the level of K-S ancestry in a group, for example higher in Tswana compared to Pedi (**Fig 3e, 3f**). Moreover, the K-S ancestry, when found to impact, seems to mainly affect *Ne* estimates older than 20 generations.

### Impact of population structure on phenotype variation and association studies

To explore the possible phenotypic implications of the fine-scale population structure, we compared allele frequencies of SNPs associated with various phenotypes (identified using the GWAS catalog) between the SEB groups. The comparison (**Fig. 4a**) shows almost six-fold variation in frequency of the *APOL1* variant rs73885319 among Sotho and Xhosa (MAF=0.03 and 0.18, respectively). Similarly, alleles in genes such as *HERC2* (associated to skin colour), *PCSK9* (associated to lipid level phenotype) and *FTO* (associated to obesity) also showed three-fold or higher allele frequency differences between SEB groups (**Fig. 4a).** A detailed list of 919 SNPs showing a minimum of three-fold difference in allele frequency is presented in **Supplementary Table 7**.

**Fig. 4.**
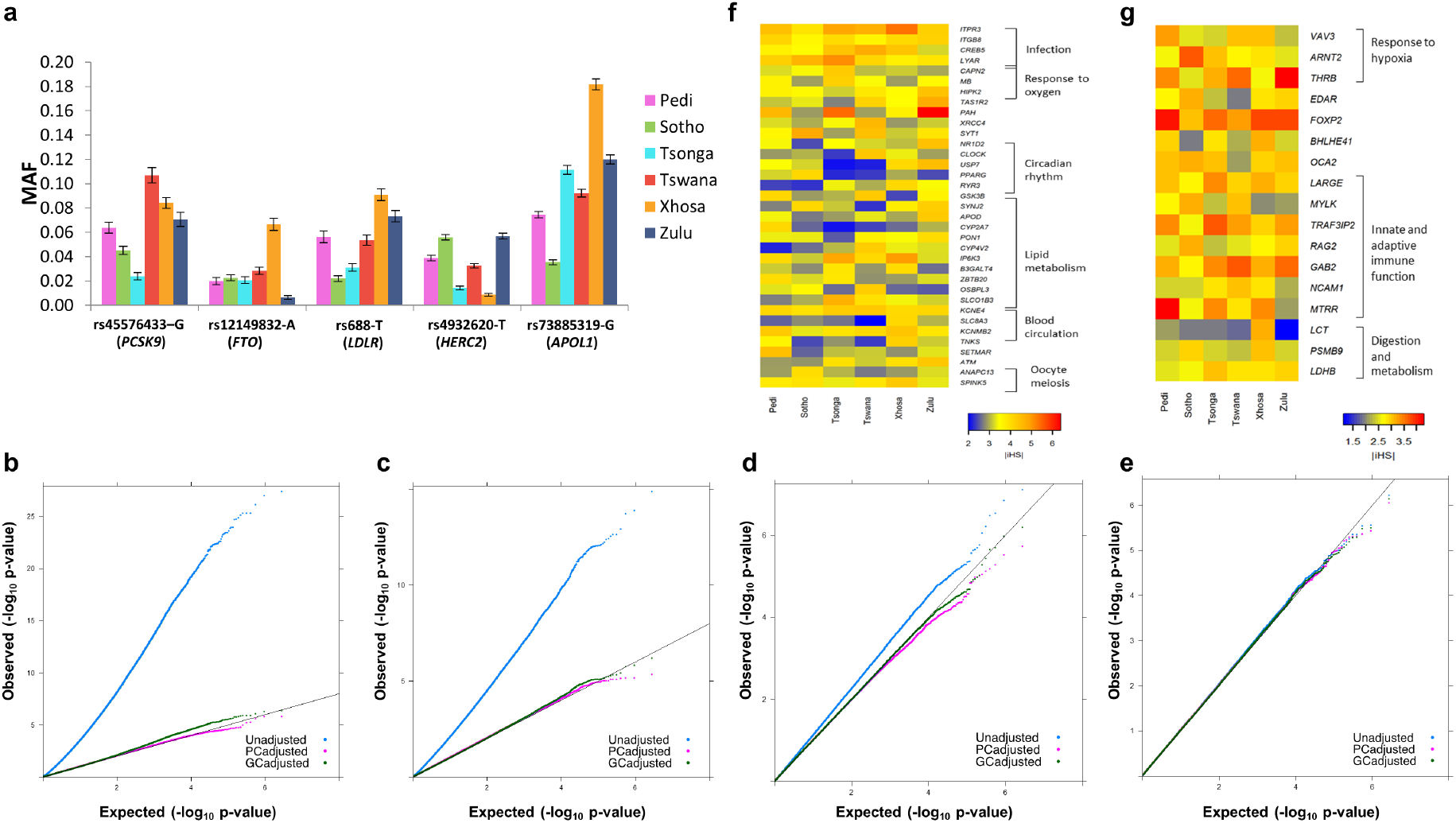
Possible impact of population structure within the South Eastern Bantu speaking (SEB) groups on genome-wide association studies (GWASs) and evolutionary estimates. **a**, Allele frequency variation of some of the well-known phenotype associated SNPs. The mean and the standard error was estimated using 50 random resampling runs. **b-e**, Representative QQ plots showing results from simulated trait GWASs comparing randomly sampled participants from (**b**) Agincourt (AGT) as cases to Soweto (SWT) as controls (**c**) 62.5% AGT+ 37.5% SWT participants as cases to 100% SWT participants as controls (**d**) Random samples from SWT (without Tswana) as cases to random samples from SWT with Tswana as controls (**e**) Randomly sampled individuals from SWT as cases and controls. For **b-e**, blue dots represent raw *P*-values, whereas green and purple dots represent *P*-values after principal component and genomic control based correction, respectively. **f**, Heatmap showing differences in iHS statistics for some of the SNPs that were detected as outliers (|iHS| >4; *P*-value<0.003) in at least two of the SEB groups. **g**, Heatmap showing differences in iHS statistics for SNPs in genes previously reported to be under positive selection, that were also detected to show moderate scores in one or more of the SEB groups (|iHS|>3, *P*-value <0.05).

Population structure accompanied by high allele frequency differences could have major implications for genome-wide association studies (GWAS). Therefore, to assess the extent to which the observed structure could bias association results, we conducted four categories of simulated traits GWASs (binary trait) using study sites (AGT, DKG and SWT) and/or ethno-linguistic labels as ‘trait-proxies’ (**see Methods**). Category 1 was aimed at stimulating a scenario where cases and controls are sampled from different study sites. The **Fig**. **4b** shows a representative QQ plot for AGT-SWT (AGT as cases, SWT as controls) comparisons, which reflects a very strong population structure with exceptionally high (>4.5) genomic inflation scores (GIS). Category 2 represents a scenario where cases are randomly drawn from two sites (AGT and SWT), while controls were from one site only (SWT). The QQ plot for this category (**Fig. 4c)** shows that even using about half of the samples from AGT could lead to substantially high (>=2) GIS and large-scale deviations. Category 3 represents the situation when both cases and controls are drawn from the same site (SWT), but have preferential representation of SEB groups. **Fig. 4d** shows that even ethno-linguistic stratification within a study site (SWT) resulted in the QQ curve reflecting population structure. Category 4 compares randomly assigned case and control status to individuals from the same site. As demonstrated in **Fig. 4e,** no major inflation was observed for this category.

The full results for 50 simulations (summarized in **Supplementary Table 8)** shows a substantial number of possible false positives associations in Categories 1 and 2, at the generally accepted genomewide *P*-value threshold of 5×10^−8^. While the genomic inflation normalizes with homogenization of the dataset, GWASs for category 3 and to a lesser extent category 4 generated false positives in a few simulations (**Supplementary Table 8**). Moreover, for each category, a substantial number of additional false-positive signals were detected at the suggestive *P*-value threshold of 5×10^−5^, some of which, with slight changes in sample sizes could easily move below the genome-wide significance threshold (**Supplementary Table 8**). We also evaluated the extent to which the two standard approaches (genomic control (GC)-based correction and PC-based correction) can control genomic inflation and possible false positives in each category^46,47^. The results (**Fig. 4b-e and Supplementary Table 8**) suggest that while both approaches are effective, in some cases they fail to remove all the genome-wide significant associations due to population structure. Therefore, linear mixed model (with PC and kinship matrix as covariates) or other advanced approaches to address the population structure^46,48^ could be more suitable for a GWAS involving SEB groups.

In order to reduce false positives due to small sample sizes, we restricted our simulations to only include common variants (MAF>0.05). The addition of rare variants (MAF<0.05) in a real GWAS, as well as increasing this dataset size by imputation, as is commonly performed in GWASs, could further increase false positives. Many of the signals from the simulated-trait GWASs have been previously reported as trait/disease genetic associations in the GWAS catalog^49^ (**Supplementary Table 9**). Therefore, in a sample set containing an unbalanced (ethno-linguistically or geographically) proportion of SEB groups, the observed associations in a GWAS could give false associations resulting from intrinsic differences between these groups, rather than an association with the trait being investigated.

### Signatures of positive selection

We used a haplotype homozygosity based selection scan to identify and compare outlier signals in the major SEB groups. The comparisons (**Fig. 4f**, **Fig. 4g, Supplementary Table 10, Supplementary Fig. 12, Supplementary Note 5)** show several of these signals (in *SYT1, PAH, CAPN2, SLC8A3* genes) to reach outlier threshold in some of the SEB groups but not in others, suggesting that the fine-scale structure can also influence evolutionary analyses. A population branch statistics (PBS) approach further detected SNPs showing high differentiation between Tswana and Tsonga (**Supplementary Table 11, Supplementary Note 5**). Many of the SNPs showing outlier PBS scores mapped to immunity related genes such as *NFKBIE, VWF* and *ITGB2* (**Supplementary Table 11)**.

### Preferential K-S gene flow

To study possible instances of preferential K-S gene flow, we identified genomic regions deviating more than ±3SD of the estimated average of K-S ancestry in each SEB group. Despite the differences in the overall K-S ancestry levels in these groups, we observed multiple genomic regions to show very high K-S ancestry in more than one of the SEB groups (**Supplementary Table 12; Supplementary Fig. 13**). For example, an extended region on chromosome 6 containing the *GRM4, HMGA1, NUDT3* genes shows high K-S ancestry in Pedi, Tsonga and Swazi, and part of this region was also observed to be K-S enriched in Venda. Similarly, another region in chromosome 6 around the *TRDN* gene shows K-S enrichment in Zulu and Xhosa. A K-S ancestry region each in Tsonga (around *DAXX* and *ITPR3* genes) and Pedi (around *ZBTB20* gene) also harboured selection outliers hinting at possible postadmixture selection scenarios (**Supplementary Table 12)**.

## Discussion

More than 40 million South Africans speak one of the 9 major South Eastern Bantu languages as their first language. Notwithstanding clear divisions in the South Eastern Bantu language phylogeny and geographic stratification of the speakers, very few studies have investigated the genetic differentiation between SEB groups. Based on a large-scale study of over 5,000 participants representing 8 of the 9 major SEB groups in South Africa, we have demonstrated the presence of a robust fine-scale population structure within the SEB groups, which broadly separates genomes of SEB groups into to the three major linguistic divisions (Nguni, Sotho-Tswana, and Tsonga), and also reflects the geographic distribution of LMAs to a large extent. The resolution of this structure within the SEB groups was enhanced considerably by taking ethno-linguistic concordance of individuals and their geographic locations into account. However, it needs to be noted that, self-identity itself is complex with about one third of the participants having more than one parent or grand-parent with a different ethnic self-identity. Moreover, while the PCA and PCA-UMAP shows clear population structure, there are exceptions highlighting the fluidity of cultural identity. Thus, self-selected group-identity encompasses significant group-related genetic variability, and it is important to emphasise that cultural identity and genetic variation are not necessarily aligned.

In alignment with results from previous studies^10,28^, our data also shows that differential K-S gene flow plays a major role in the population structure of SEB groups. However, the persistence of the structure even after accounting for differential K-S admixture suggests the contribution of other demographic factors in the genetic differentiation of these groups. The SEB groups start to show clear divergence in population size dynamics from about 40 generations ago. This time frame converges with the earliest dates of K-S admixture and probably points at the initiation of migration events that gradually separated these groups. On the other hand, a rather wide variation in K-S admixture dates (spanning ∼20 generations) among SEB groups possibly reflects the complexity of the settlement of different parts of the country by the ancestral BS populations. Comparison of present-day SEB groups with Iron-Age farmer genomes provided evidence for genetic continuity in a geographic region in Central-East South Africa for at least the last 300-500 years. Our results, while attesting to the well-known pattern of K-S female-biased gene flow, showed notable differences in the extent of this bias among different SEB groups demonstrating that the nature of interaction between K-S and BS could have varied temporally and geographically.

The dataset we generated for this study has provided a much better contextualization for previously sequenced iron-age genomes from Southern Africa. The SEB are unique in Africa, as being among the very few populations that contain considerable gene flow from the Khoe-San. These data therefore are of major importance in terms of understanding the interaction between the Khoe-San and other Southern African populations. They will play an important role in providing insights through comparative analyses once more genetic data from hunter-gatherers and ancient genomes from this geographic region become available.

Our analyses including allele frequency comparisons, genome-wide scans for selection and K-S ancestry distribution show the SEB groups to be highly diverged at certain genomic regions. Based on simulated trait GWAS, we further illustrate that the fine-scale population structure within the SEB groups could impact a GWAS by introducing a large number of false positives. A combination of cautious study design to minimize geographic and ethno-linguistic biases and stringent measures for population structure correction is therefore recommended for GWASs involving SEB groups. Moreover, while GWAS can address the false positives introduced due to population structure using GC, PC or other approaches, it is impossible to identify and control for population structure in candidate gene studies. Therefore, utmost care should be taken during study design to ethnically and geographically homogenise samples in order to control for false positives in association studies using limited markers.

A major limitation of our study is that the sampling sites do not cover the full geographic spread of SEB groups in the country, possibly causing some of the groups to be sub-optimally represented in our dataset. Nevertheless, our results suggest that we are at a critical point in history where the population structure is still observable with efficient sampling and in-depth ethno-linguistic characterization, even if it is gradually diminishing due to migration and intermingling between different SEB groups. We hope that our findings will motivate studies with larger sample sizes and wider geographic representation to help unravel the demographic events that contributed to the peopling of South Africa.

## Data availability

The data is being submitted to the European Genome-phenome Archive (EGA, http://www.ebi.ac.uk/ega/) (we will update the EGA ID once it becomes live) and can be accessed through the H3Africa-DBAC.

## Code Availability

All software and analysis code is publicly available The code for *plotY is* is available through GitHub (https://github.com/shaze/ymthaplotools).

## Ethics

This study was approved by the Human Research Ethics Committee (Medical) of the University of the Witwatersrand (Wits) (protocol number M121029), and renewed in 2017 (protocol number M170880). In addition, research at the Dikgale Study Centre was approved by the Medunsa Research and Ethics Committee of the University of Limpopo (MREC/HS/195/2014:CR). Community engagement preceded sample collection and all participants provided broad consent for medical and population genetic studies.

## Acknowledgements

We wish to express our profound gratitude to the more than 5000 unnamed participants who took part in the study and provided blood samples, as well as the team of field workers, laboratory scientists and administrators who made the sample collection possible. We would like to acknowledge Adrian Frith for providing us with the coordinates of linguistic majority areas of each SEB group and Prof Christopher Mathew for helpful comments and feedbacks about the manuscript.

## Author contributions

Study design: DS, AC, CS, MR, and SH

Data preparation and initial processing: SH

Analysis: DS, CFL, AC, SA, SH

Archaeological, linguistic and historical data interpretation: GW, HG, NC-P, KB

Writing: DS and AC with contributions from all other authors

Other contributions: ST, FG-OC, LM, SN, FM and MA directed the field work and sample collection

All authors critically evaluated and approved the manuscript.

## METHODS

### Sampling and genotyping procedures

The volunteers included in this study were sampled across three study sites (**Fig. 1a**); Agincourt (AGT), Dikgale (DKG) and Soweto (SWT) under the Africa-Wits-INDEPTH partnership for genomic studies (AWI-Gen) project as part of the Human Heredity and Health in Africa (H3Africa) Consortium^50^. Of these SWT is urban, whereas DKG and AGT are rural/semi-urban sites. The study included a total of 5,268 individuals (mostly within the age range of 40 to 60 years) representing 8 major South African SEB groups: Tsonga, Pedi, Venda, Sotho, Tswana, Swazi, Zulu, and Xhosa. Details of community engagement, written informed consent, and genomic DNA extraction from blood samples have been described elsewhere^51^. The samples were genotyped on the H3Africa array (∼2.3M SNPs) using the Illumina FastTrack Sequencing Service2. The default Illumina pipeline was used for the genotype calling (build GRCh37/hg19).

### Data quality control procedures

Quality control (QC) on the AWI-Gen genotype dataset was performed using PLINK (v1.9)^52^ and involved removal of duplicate SNPs, multi-allelic SNPs, INDELs and SNPs with a missingness >0.05, MAF <0.01 and SNPs that failed HWE test (*P*-value <0.0001). Individuals with missingness >0.05, discordant sex information and lacking self-reported ethnicity information were also removed. The genotype dataset post-QC consists of 5,056 samples and 1,733,001 autosomal SNPs (AWI-S1) (**Table 1** and **Supplementary Table 13**). A linkage disequilibrium (LD)-pruned version of this dataset was generated by removing SNPs in high LD (*r^2^*>0.5 within a window of 50 SNPs, and with a window slide of 5 SNPs) using PLINK. The same parameters for LD-pruning were used for the datasets described below.

### Assessment of Relatedness

To identify related individuals, we estimated identity-by-descent (IBD) segments for each sample pair, based on the LD-pruned AWI-S1 dataset. For each pair of related individuals (PIHAT >0.18), the sample with higher missingness was dropped, resulting in the removal of 737 SEB participants in the process leading to AWI-S2 dataset (**Table 1**). We also estimated genetic relatedness for all pairs of individuals in the AWI-S2 dataset using KING^53^ and GENESIS^54^ and PC-Relate option for plotting. After these QC-steps, no first-degree or second-degree relatives were found in the dataset used for the analyses below (**Supplementary Fig. 14**).

### Analysis of ethno-linguistic concordance

In addition to self-reported ethnicity of the participant, the study also captured self-reported ethnicities of the parents and grandparents of each participant. Admixture within South-Eastern Bantu-speaking (SEB) groups as well as between SEB and non-SEB groups has been common in recent South African history. Since admixture events could influence fine-scale comparisons between SEB groups, we identified the participants that were ethno-linguistically concordant (EC), i.e. have reported the same ethnicity for themselves, both parents and the four grandparents (allowing for a maximum of one mismatch). This set of 2702 EC participants was defined as AWI-S3 dataset, details are listed in **Table 1**).

### Sample size homogenisation

The representation of various SEB groups in the AWI-Gen study was notably skewed toward Tsonga, Pedi and Zulu (with over 2,000, 1,200 and 600 samples, respectively) (**Table 1**). To avoid bias due to sample size differences and make the population sizes of the SEB groups comparable, we randomly downsized these three large groups to 80 individuals for each group from the AWI-S3 dataset. This dataset referred to as AWI-S4 consists of 484 samples (80 Pedi, 46 Sotho, 33 Swazi, 80 Tsonga, 73 Tswana, 29 Venda, 63 Xhosa, and 80 Zulu individuals) (**Supplementary Table 13**).

### Data merging workflow

For comparison of our population with previously studied populations, the AWI-S2 data (4319 SEB unrelated samples) was merged with additional world-wide datasets from (ref. ^21^), 1000 Genomes Project (KGP)^55^, and African Genome Variation Project (AGVP)^28^ using PLINK (hereafter Merged dataset 1), and only the SNPs that overlapped between all datasets were retained (**Supplementary Table 13**). We also generated another dataset (hereafter Merged dataset 2) by merging the abovementioned dataset with data from Bantu-speaking groups in South Africa, e.g. the Southern African Human Genome Project (SAHGP)^27^, and Mozambique^31^ (**Supplementary Table 13**). In addition, the AWI-S3 was also merged with four Iron-Age samples with Bantu-related ancestry presented in ancient DNA studies^10,11^. An additional dataset based on merging K-S data^56^ to AWI-S3 was generated for X chromosome analysis. This layered merging was performed to retain the maximum number of SNPs possible for each analysis.

### Exploring population structure

To investigate the population structure within the SEB groups, principal component analysis (PCA) was performed on the basis of the LD-pruned AWI-S2 dataset using the program smartPCA implemented in the EIGENSOFT suite^57^. Additionally, PCA was also performed first on the basis of the LD-pruned AWI-S3 dataset, and then for the Merged dataset 2. To further investigate the population structure obtained in PCA results, we combined the information for the first 10 PCs using a non-linear dimensionality reduction tool, called uniform manifold approximation and projection (UMAP)^30^.

### Genetic distance between SEB groups

To investigate genetic affinities between the different SEB groups, we estimated Weir and Cockerham’s F_ST_ statistics (Weir and Cockerham 1984) between pairwise SEB populations included in the Merged dataset 2 (EC-downsized) using PLINK. The relationship between the SEB groups based on pairwise F_ST_ value was represented with a UPGMA tree using the program MEGA X^58^.

### Linguistic phylogeny of SEB languages

The linguistic phylogeny is based on lexical data for 100 concepts in 69 Bantu varieties, 34 of them part of South-Eastern Bantu languages and 35 outgroup languages belonging to different major Bantu branches^59^. The lexical data were binary recorded in 1304 partial cognate sets (form-meaning associations). The resulting matrix was analysed with Bayesian inference methods as implemented in MrBayes (v3.2.7)^60^ using a restriction-site model^61^.

### Correlations between geographic and genetic distance

Mantel tests were implemented to investigate possible relationships between the geographic and genetic distances between the SEB groups. As many of the groups such as Zulu and Xhosa were sampled from sites that are quite distant to their native geography, we calculated geometric medians of the population of speakers for each language using Weiszfeld’s algorithm (http://www.or.uni-bonn.de/~vygen/files/fl.pdf), and considered them as the midpoints of each group. The great circle geographic distance between each midpoint was estimated using an online tool (https://www.geodatasource.com/distance-calculator). The genetic distance matrix was based on weighted mean F_ST_ estimates for each pair of SEB groups. The Mantel test was performed using the R package *vegan^62^*, using 9999 permutations to test the correlations between the geographic distances and F_ST_ based genetic distances.

### Estimating admixture dynamics

For global ancestry inference, we used an unsupervised clustering algorithm implemented in ADMIXTURE (v1.3)^32^ on the Merged dataset 2 (EC-downsized). The number of K-groups analysed varied from *K*=3 to *K*=8, and 50 independent runs with a random seed for each K-group was performed. The K-group with the lowest cross-validation error (CV) was considered “optimal”. PONG^63^ was used for merging and visualizing the clustering outputs of all the runs from the ADMIXTURE analysis, and major modes were used for the ADMIXTURE plots. To compare the differential contributions of the main ancestries (K-S, BS and Eurasian component), the average admixture proportions of each ancestry was computed from the ADMIXTURE results at *K*=3 for each SEB group in the Merged dataset 1. For each study site, we further estimated the average admixture proportion for each ancestry at *K*=3 in each SEB groups. We applied a t-test to compare whether there are significant differences in K-S ancestry proportion across the three sites for a given SEB group.

### Local ancestry deconvolution

For local ancestry inference, we used RFMix (v1.54)^64^, on the basis of the Merged dataset 1 (EC). As reference panels, we selected: YRI for Central-West African ancestry; CEU for Eurasian ancestry; and combined Ju|’hoansi, /Gui //Gana, and Karretjie^21^ for K-S ancestry. The merged dataset was first phased using SHAPEIT2^65^ with a reference panel of worldwide haplotypes^55^, and then analysed with two runs of expectation maximization (EM=2), forward-backward and PopPhased options. The genetic map from HapMap Phase 2 build GRCh37/hg19 was used for the analysis.

### Ancestry specific PCA

To investigate whether the differential K-S gene flow is the only factor leading to the observed population structure, SEB haploid genomes were masked for regions of K-S and European ancestries identified using RFMix. We then analysed haploid regions with more than 50% Bantu-related ancestry using the ancestry-specific PCA (AS-PCA) approach^66^.

### Admixture date inference

To reconstruct the timeframe of admixture events between the major ancestry components in SEB populations, we used three admixture dating methods, fastGLOBETROTTER, the recent implementation of GLOBETROTTER^33^, MALDER (v1.0)^38^ and MOSAIC (v1.3.7)^39^. The details for each method is described in **Supplementary Note 2**.

### Comparison with Iron-Age genomes

To compare the genetic affinities of modern SEB groups to Iron-age Bantu-related samples from Southern Africa, we analysed the AWI-S3 dataset together with five ancient samples: four associated with Iron-Age (300 to 500 year old) farmers in South Africa^10^, and one 700 years old sample from Pemba, Tanzania^11^. We used smartPCA to project the ancient samples onto the modern samples (using the following options: lsqproject=YES; killr2=YES; and shrinkmode=YES). To better visualize genetic affinities between ancient and modern samples, we performed the PCA-UMAP analysis using the PC coordinates for the first ten PCs and UMAP tool for the analysis ^30^, and a custom Python script for the plotting.

### Y chromosome and mitochondrial haplogroup analysis

Y-haplogroup analysis was carried out using our new in-house *plotY* tool (https://github.com/shaze/ymthaplotools), based on a modified version of the tree and mutations table of AMY-tree^67^. The results were then validated using SNAPPY^68^. MtDNA haplotyping was performed using Haplogrep 2^69^, using Phylotree mtDNA tree build 17rsrs-RSRS^70^. The details for mtDNA and Y-haplogroup detection are described in **Supplementary Note 3**.

### Sex-biased admixture patterns

Recent literature has suggested the comparison of X-chromosome and autosomal contributions from the two source population as a robust method to test for possible sex-biased admixture^71^. To investigate the extent of sex-bias in the contributions of different ancestral populations to admixed SEB groups, the AWI-S3 and YRI and CEU from KGP were merged with available data^56^, consisting of the 33 K-S samples. The ancestry proportions for each were estimated using ADMIXTURE at *K*=3 for three datasets: the X-chromosome dataset, the autosomal dataset, and the merged autosome-X-chromosome dataset. Admixture difference (ΔAdmix) ratios were then calculated using the method proposed by (ref. ^72^). A positive ΔAdmix ratio indicates an excess of female-specific admixture contributions, while a negative value indicates an excess of male-specific admixture. To test statistical significance of the difference between the ΔAdmix for each ancestry between pairs of populations, we used the Wilcoxon rank-sum test.

### Population size dynamics

To estimate and compare the variation in recent population size of the different SEB groups, IBD segments were detected from the downsized Merged dataset 1 using the program IBDseq^73^ for each group (with default parameters). The output was then used as input for the program IBDNe^43^, which computes the effective population size (*Ne*) for each SEB group for the last few hundred generations. To avoid the conflation effect of short IBD segments^74^, only IBD segments longer than 4cM were retained for the *Ne* estimation, and the remaining parameters were set as default.

Ancestry-specific effective population size (AS-IBDNe)^43^ was estimated for the different SEB groups to identify the contribution of K-S and BS ancestries to the *Ne* dynamics of various SEB groups. This analysis was performed on the dataset that was used to estimate the overall *Ne*. We followed the pipeline provided by the authors, which implements both IBD and local ancestry information from the genotype data. The first step in this approach was to phase the data using Beagle (v5)^75^, and then IBD segments were detected using RefinedIBD and local ancestry information was inferred with RFMix (YRI, K-S and CEU were used as reference source populations). Finally, IBDNe was used to estimate the ancestry specific *Ne* from the detected IBD segments and the ancestry blocks inferred from the local ancestry analysis.

### Allele frequency variation of phenotype associated variants

We used PLINK to estimate allele frequencies of all SNPs in our dataset that are included in the GWAS catalog^49^ (accessed on 19^th^ April 2020), in the six major SEB groups (represented by at least 80 individuals in the AWI-S4 dataset). Standard error for allele frequencies was estimated using 50 bootstrap iterations in a subset of 30 individuals from each SEB group.

### Simulated genetic associations to illustrate the potential effect of population structure

To simulate various possible scenarios for genetic association studies using ethno-linguistically and geographically mixed set of SEB participants, four categories of artificial “case-control” trait simulations were performed. In the first category, the sampling site was used as the basis for assigning the case and control status. Here, the “case” label was assigned to 800 randomly sampled individuals from one of the three sites and the “control” label assigned to 800 randomly sampled individuals from a different site. Independent comparisons, 50 iterations each for AGT-DKG; DKG-SWT; AGT-SWT were performed. The second category corresponds to a scenario in which cases (n=800) are a mixture of samples from AGT and SWT and the controls (n=800) are sampled from SWT only. Three sets of cases with varying proportions of AGT and SWT representation (37.5% AGT+ 62.5% SWT, 50% AGT +50% SWT and 62.5% AGT +37.5% SWT) were generated and 50 iterations were performed for each set. The third category of trait simulation was aimed at studying the impact of ethno-linguistic stratification within a sampling site, SWT. For two sets (50 iterations, 500 cases-500 controls) generated in this category, the assignment was done in a way in which one of the ethno-linguistic groups (Tswana in set 1 and Tsonga in set 2) was absent in cases but present in controls. The fourth category, was generated by randomly assigning case and control labels to the samples form a single site at a time. GWAS for each of the case-control pairs in all the sets under the four categories were conducted using the association testing function in PLINK. Genomic inflation scores were recorded for each run, and signals at a genome-wide significance threshold of *P*-value=5×10^−8^ were identified, as well as a less stringent suggestive significance threshold (*P*-value=1×10^−5^). To assess the extent of population structure correction possible with a genomic control based approach, for each run the inbuild correction testing function was implemented using the “adjust” flag in PLINK. To assess the impact of PC based correction, for each of the case-control iterations, PCs for the dataset was estimated using PLINK and the first three principal components were used as covariates in logistic regression based association testing in PLINK. QQ plots were generated using a custom R scripts. Possible phenotypic roles of the associations detected in these simulated trait GWASs were assessed using the GWAS catalog^49^.

### Genome-wide scans for selection

To identify SNPs under positive selection, we calculated the integrated haplotype homozygosity scores (iHS) ^76^ implemented in the program *Selscan* (Szpiech and Hernandez, 2014). The AWI-S4 dataset was used for this analysis, and only SNPs with MAF <0.05 were considered. We included six SEB groups, and we removed Venda and Swazi samples due to their small sample size. For each SEB group, the raw iHS were normalized across 40 frequency bins. A random sampling of scores across populations was performed to assess *P*-values for various score cut-offs. Based on this |iHS|>4 was considered as outliers (*P*-value<0.003). The mapping of SNPs to genes was performed based on information retrieved from Ensembl Biomart (Ensembl genes version 100; https://grch37.ensembl.org/biomart/).

We also used the population branch statistics (PBS) analysis^77^ to identify SNPs under positive selection. PBS is a summary statistic that utilizes pairwise Fst values among three populations to quantify genetic differentiation along each branch of their corresponding three-population tree. Since the overall genetic distance between the SEB groups is not very high, we considered only two groups from our study: the one with the highest K-S ancestry (Tswana), and the other with the lowest K-S ancestry (Tsonga). CHB from KGP was selected as the outlier population for this study. For each exonic SNP (identified using Ensembl Biomart as mentioned above) with MAF>0.01, F_ST_ values were estimated between the three pairs of the populations (CHB-Tswana, CHB-Tsonga, and Tswana-Tsonga) using VCFtools^78^. PBS scores were then estimated in Tswana-Tsonga-CHB and Tsonga-Tswana-CHB comparisons using the method described in (ref. ^77^).

### Preferential K-S gene flow

To identify genomic regions showing enrichment of K-S ancestry in the SEB groups, local ancestry inference was performed using RFMIX as described above. To avoid statistical noise, regions around centromeres and telomeres (2Mb from each side) for each chromosome were excluded from the analysis. Only SNPs with a high confidence value for K-S ancestry i.e. posterior probabilities value>0.8 were retained for the analysis. Ancestry regions (containing at least 3 SNPs) exceeding the average genome-wide K-S ancestry estimate by at least +3SDs were considered as candidates for preferential K-S gene flow. We then investigated if there are regions of adaptive introgression in the genomes by overlapping the regions under positive selection (as described above) and regions showing K-S enrichment.

