## Supplementary Figure for "Genetic-substructure and complex demographic history of South African Bantu speakers"

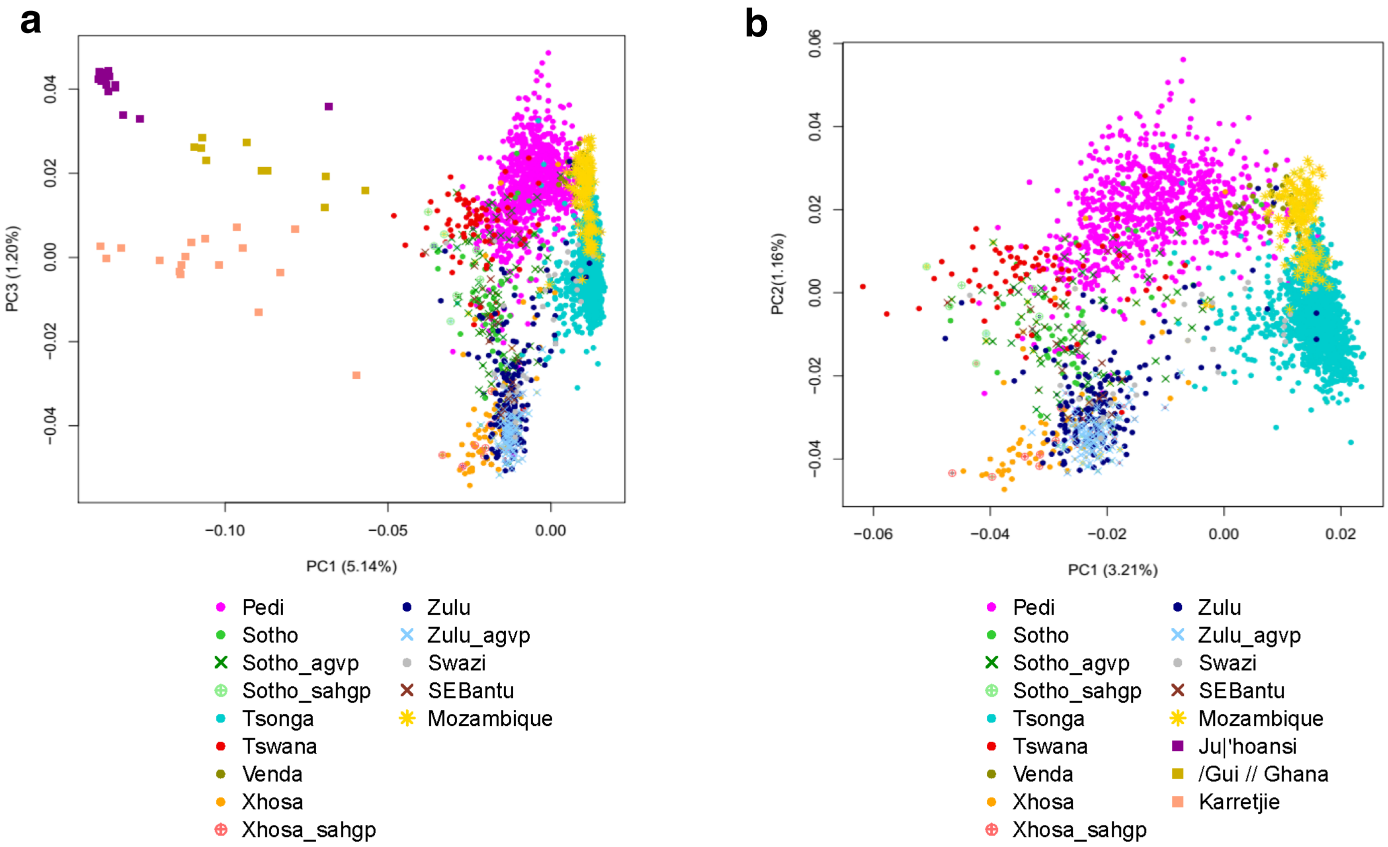


**Supplementary Figure 1**. **Principal Component Analysis (PCA) based comparison of South Eastern Bantu-speaking (SEB) groups from the AWI-Gen study to previously studied populations from Southern Africa. a,** PCA showing three Khoe-San (K-S) groups (ref. [11]) and SEB groups from: ref. [11] (named SEBantu), ref. [9] (indicated by the suffix "_agvp" in group name), ref. [18] (indicated by the suffix "_sahgp" in group name), and ref. [22] (named Mozambique). PC1 splits the K-S from SEB groups while PC3 splits the geographically southern K-S group (Karretjie People) from more Northern K-S groups as well as show the Sotho-Tswana speakers on one extreme and Nguni speakers on the other. **b,** PCA comparing SEB groups only, shows an overall concordance in localization of SEB groups from different datasets. For example, the Sotho from ref. [18] (Sotho_sahgp) and ref. [9] (Sotho_agvp) studies cluster with AWI-Gen Sotho (Sotho).

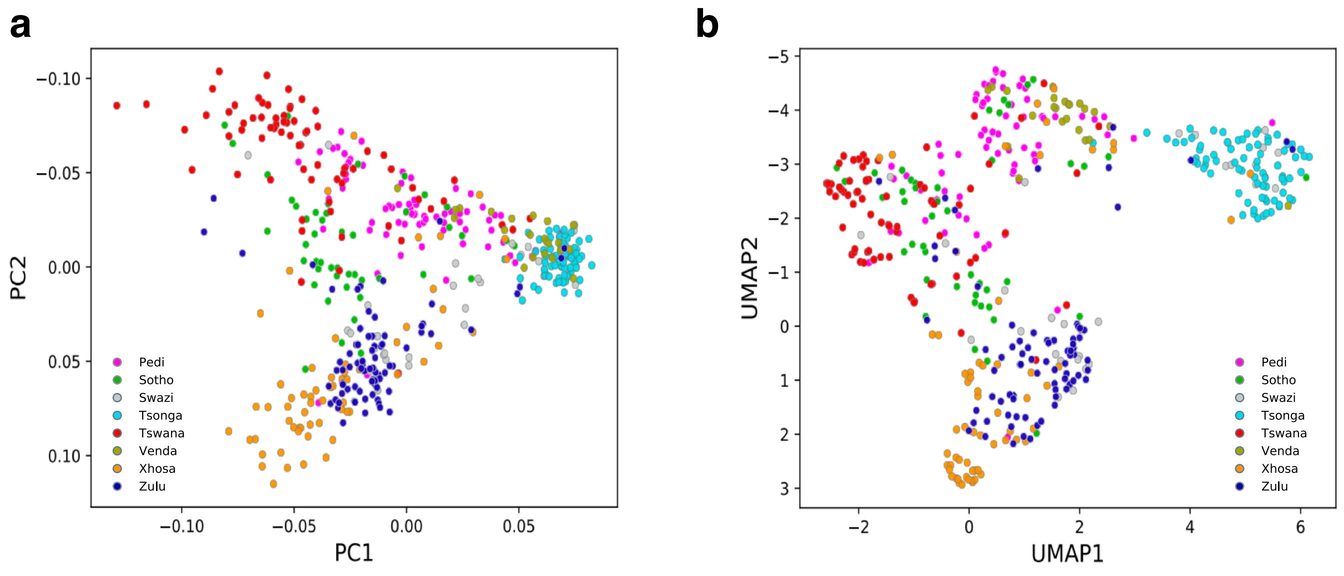


**Supplementary Figure 2. Plots representing major South Eastern Bantu-speaking (SEB) groups included in this study downsized to a maximum of 80 ethno-linguistically concordant samples for each group. a,** PC1 and PC2. **b,** PCA-UMAP plot summarizing the composite of first 10 PCs.

a

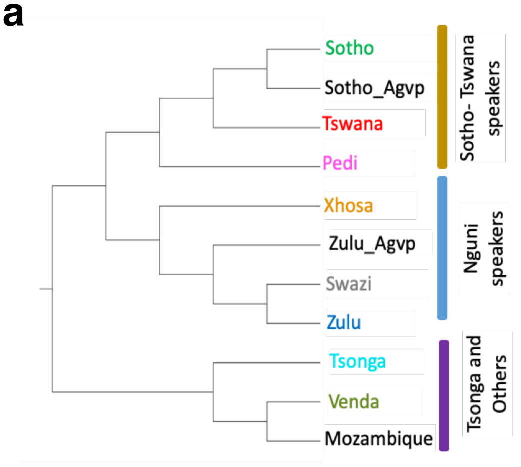


b

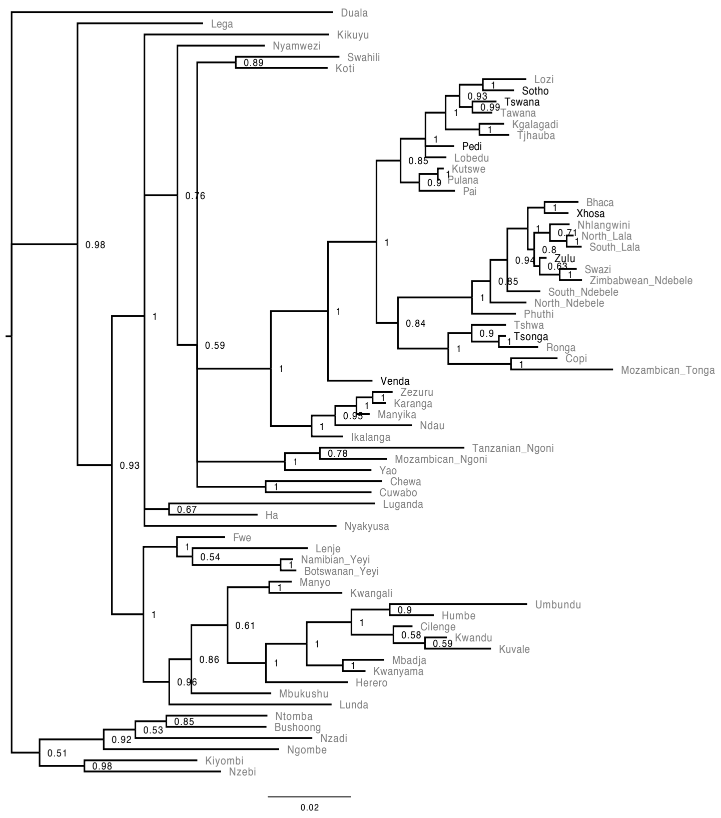


c

| SEB group | Pedi | Sotho | Swazi | Tsonga | Tswana | Venda | Xhosa | Zulu |
|---|---|---|---|---|---|---|---|---|
| Pedi | | | | | | | | |
| Sotho | 0.0004 | | | | | | | |
| Swazi | 0.0006 | 0.0004 | | | | | | |
| Tsonga | 0.0018 | 0.0024 | 0.0011 | | | | | |
| Tswana | 0.0011 | 0.0004 | 0.0021 | 0.0046 | | | | |
| Venda | 0.0015 | 0.0016 | 0.0015 | 0.0011 | 0.0044 | | | |
| Xhosa | 0.0018 | 0.0007 | 0.0012 | 0.0039 | 0.0019 | 0.0032 | | |
| Zulu | 0.0010 | 0.0005 | 0.0004 | 0.0026 | 0.0019 | 0.0027 | 0.0005 | |

**Supplementary Figure 3. Comparison of trees based on genetic and linguistic distances between the South Eastern Bantu-speaking (SEB) groups. a,** UPGMA tree for pairwise $F_{ST}$ distances between SEB groups from AWI-Gen study, ref. [9] (indicated by the suffix "_Agvp") and ref. [22] (named "Mozambique"). **b,** The full majority-rule consensus tree based on lexical data for 100 concepts in 69 Bantu language varieties, 34 of them part of South Eastern Bantu langugages. **c,** Pairwise mean $F_{ST}$ values between major SEB groups from the current study.

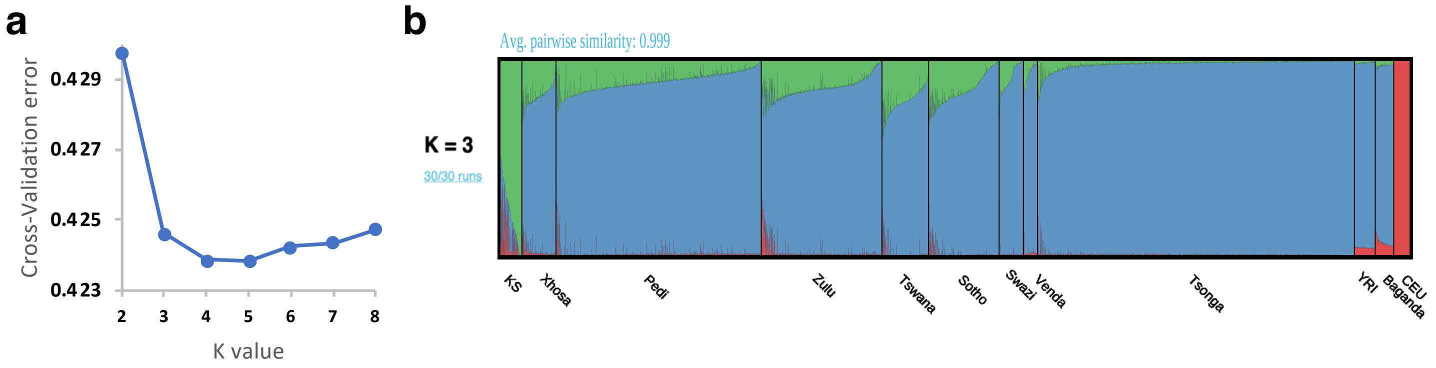


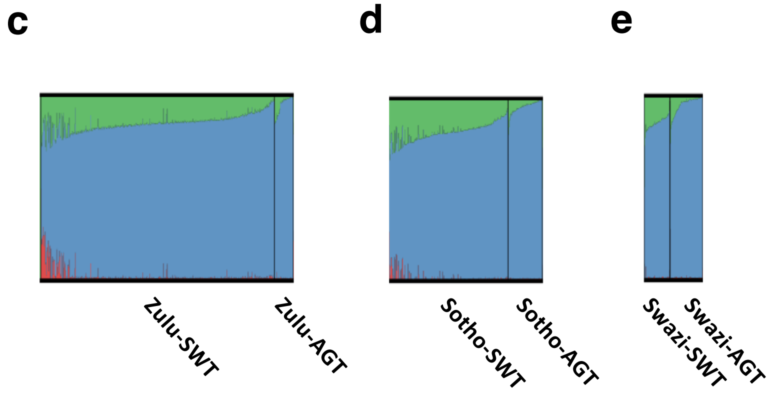


**f**

**KS ancestry difference -P value**

| | AGT-SWT | AGT-DKG | DKG-SWT |
|---|---|---|---|
| **Pedi** | 6.52E-13 | 5.80E-08 | 1.35E-19 |
| **Tsonga** | 1.29E-07 | 3.54E-07 | 3.39E-01 |
| **Sotho** | 2.13E-70 | - | - |
| **Swazi** | 1.45E-19 | - | - |
| **Venda** | - | - | 7.20E-02 |
| **Zulu** | 2.19E-42 | - | - |

**Supplementary Figure 4. Geography strongly influences the levels of Khoe-San (K-S) ancestry in some of the South Eastern Bantu-speaking (SEB) groups. a**, Cross validation value plot for ADMIXTURE plot in Figure 2a. **b**, ADMIXTURE plot at *K*=3 showing varying levels of Bantu-speaker like (blue), Khoe-San (green), and European-like (red) ancestry in all unrelated SEB individuals (n=4,319). **c-e**, shows ADMIXTURE plots at *K*=3 for three of the SEB groups- (c) Zulu (d) Sotho (e) Swazi with sampling site information (AGT- Agincourt; DKG – Dikgale and SWT- Soweto) appended to ethnolinguistic labels in the legend. **f,** *P*-values for differences in K-S ancestry levels of SEB participants from the three sites. Comparisons showing *P*-values <0.05 are shown in green. Comparisons where there are no or very few samples in one of the sites are shown with "-".

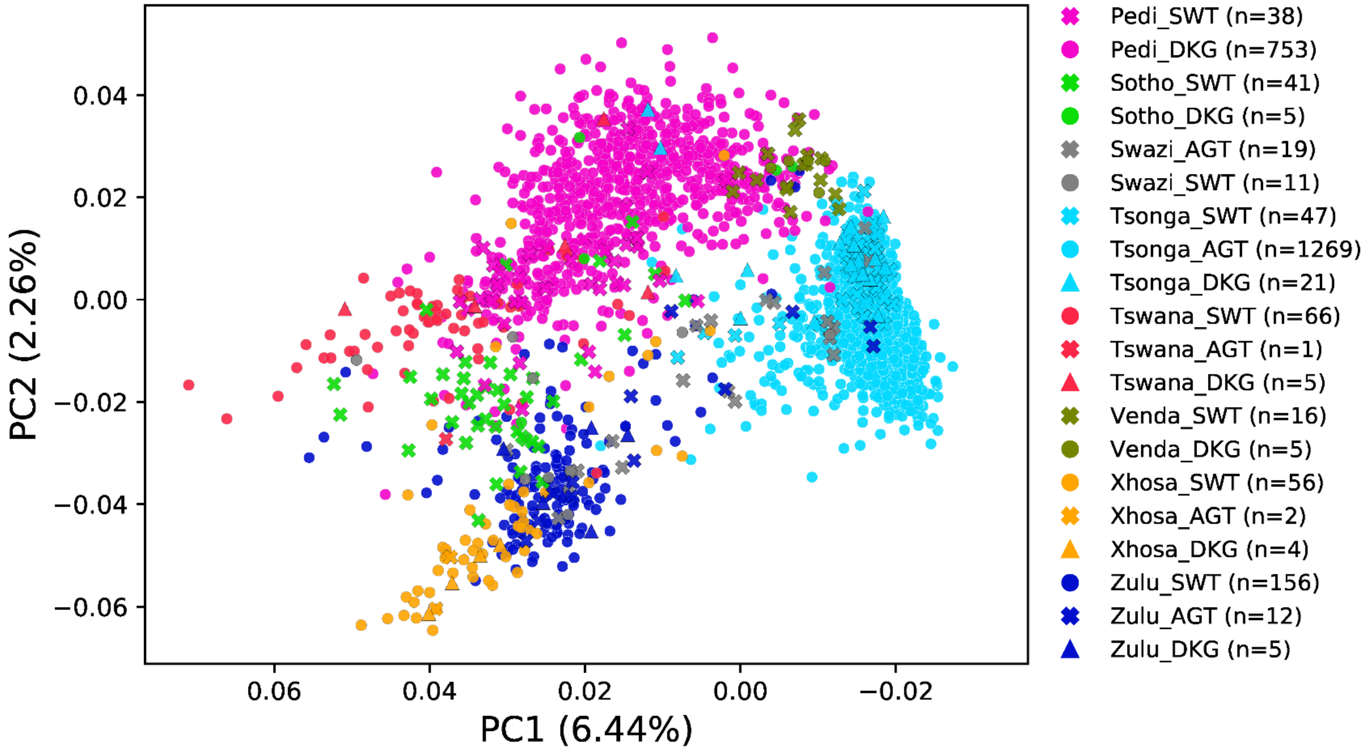


**Supplementary Figure 5. Principal Component (PC) plot showing South Eastern Bantu-speaking (SEB) groups from AWI-Gen labelled by both ethnicity and site (Soweto- SWT, Dikgale- DKG and Agincourt- AGT) of collection.** In some cases the participants instead of clustering with other members of the same SEB group from another sampling site, tend to cluster with the participants of a different SEB group sharing the sampling-site. For example, some of the Zulu and Swazi participants sampled from AGT, cluster with Tsonga (predominant group from AGT), highlighting the importance of site of sample collection.

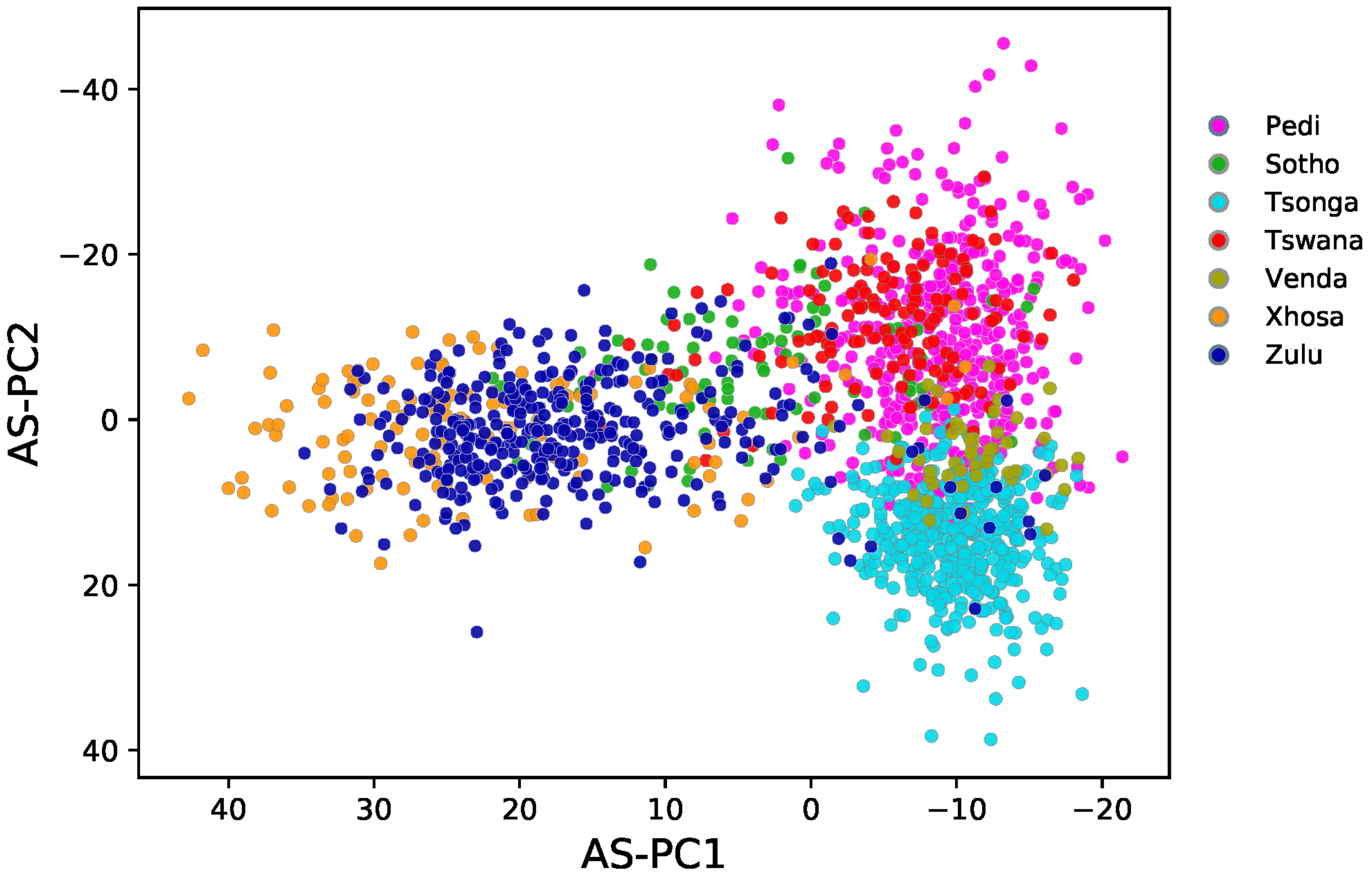


**Supplementary Figure 6. Principal component analysis (PCA) plots showing persistence of population structure in South Eastern Bantu-speaking (SEB) groups post Khoe-San (K-S) ancestry masking using AS-PCA approach.**

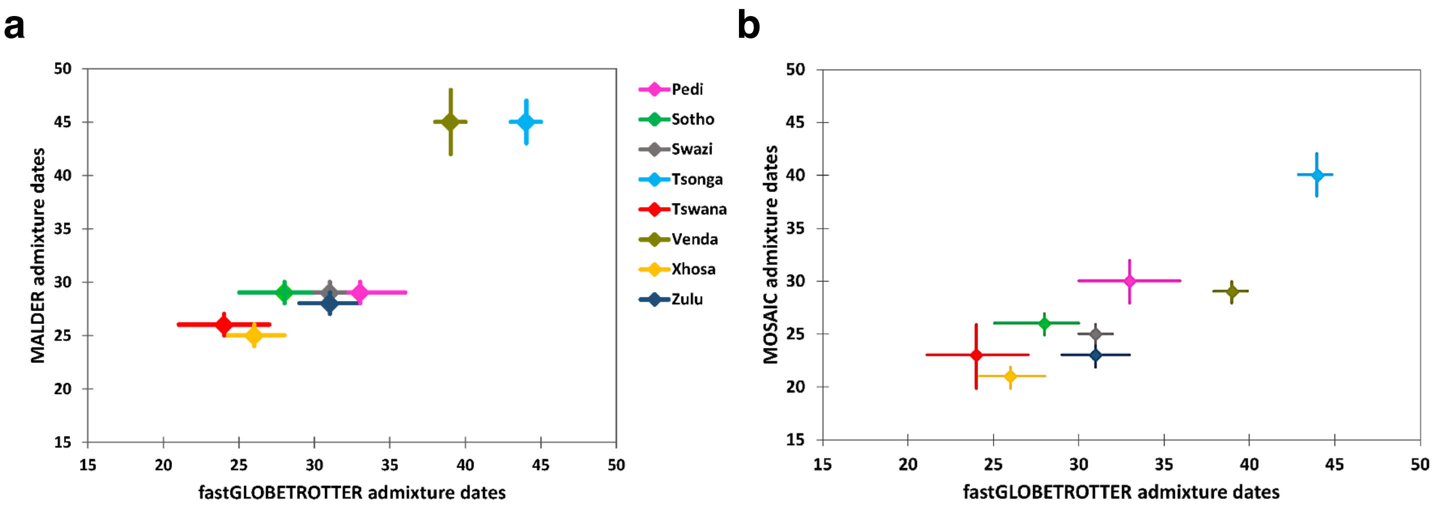


**Supplementary Figure 7. Plot showing inferred two-way admixture dates (in generations) for each South Eastern Bantu-speaking (SEB) group estimated using fastGLOBETROTTER compared to dates estimated using a, MALDER and b, MOSAIC**. Figure also showing 95% CI bars from each method obtained using bootstrapping. Further details about these dates were included in **Supplementary Table 2**.

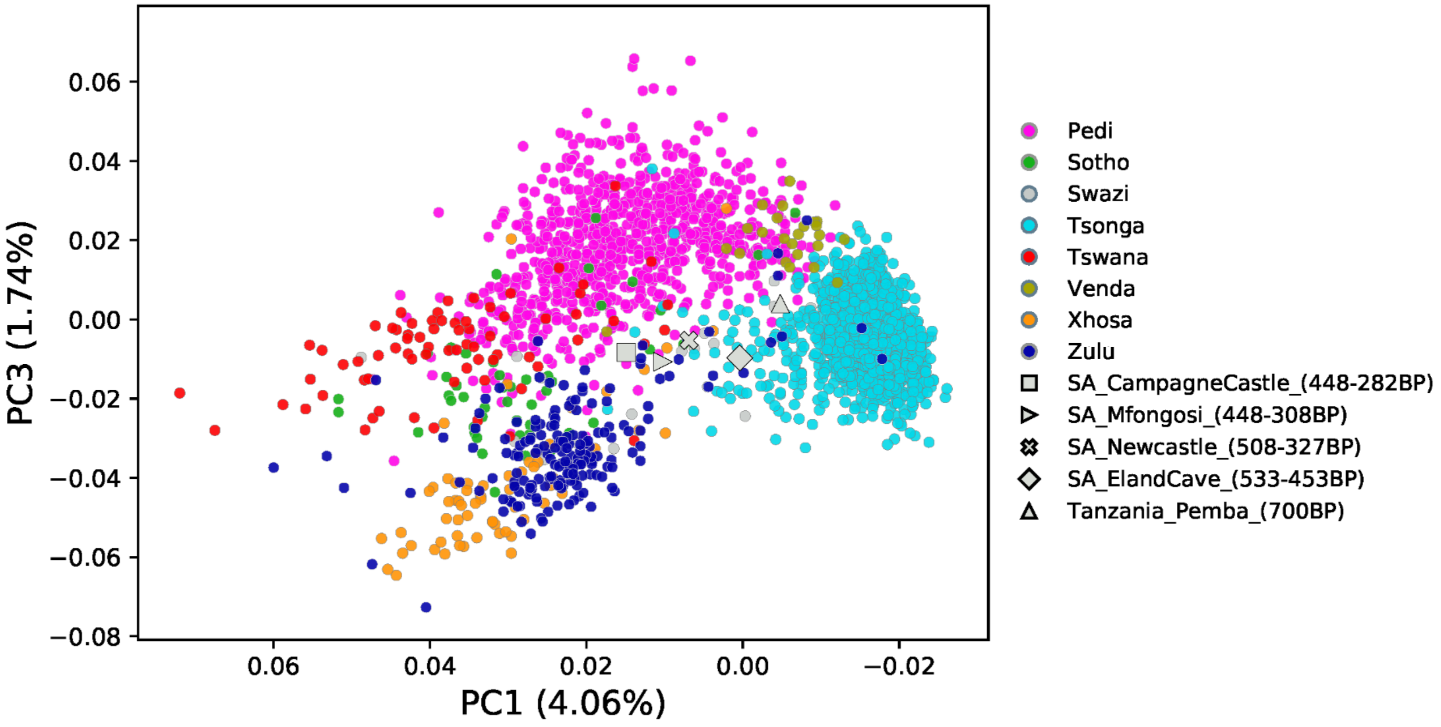


**Supplementary Figure 8. Principal component analysis (PCA) plot showing Iron-Age genomes along with South Eastern Bantu-speaking (SEB) groups from the current study.**

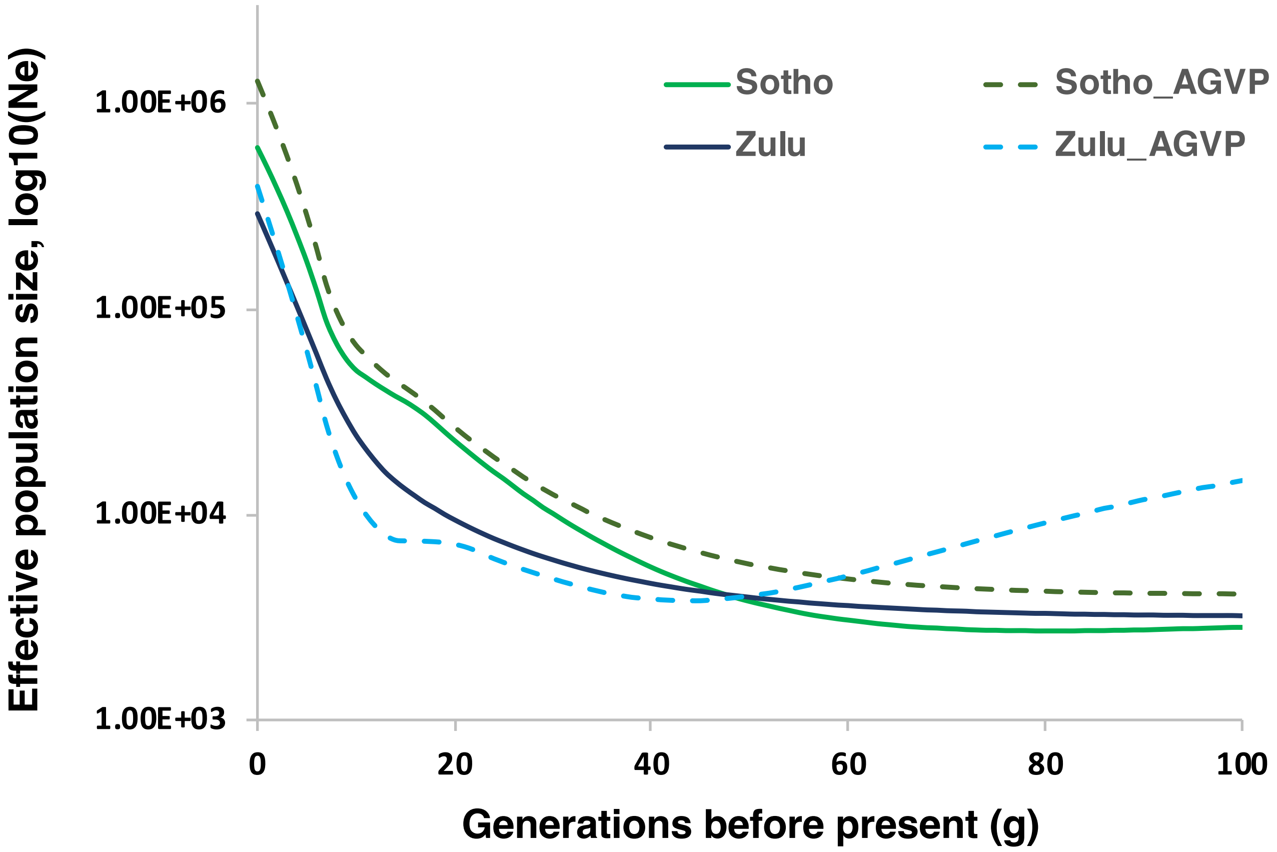


**Supplementary Figure 9**. **Comparison of effective population size (*Ne*) estimates for Sotho and Zulu from AGVP (ref. [9]) and the AWI-Gen study**. For both the datasets, Zulu maintain a lower *Ne* than Sotho around the period of ~15-40 generations. The similarity observed between the *Ne* profiles of Zulu AWI-Gen and Zulu_AGVP as well as Sotho AWI-Gen and Sotho-AGVP is despite the unequal sample size used for the analysis. For the AWI-Gen dataset, the sample size for both the groups was around ~220; while for the groups in the AGVP dataset the sample sizes were n=86 for Sotho and n=100 for Zulu.

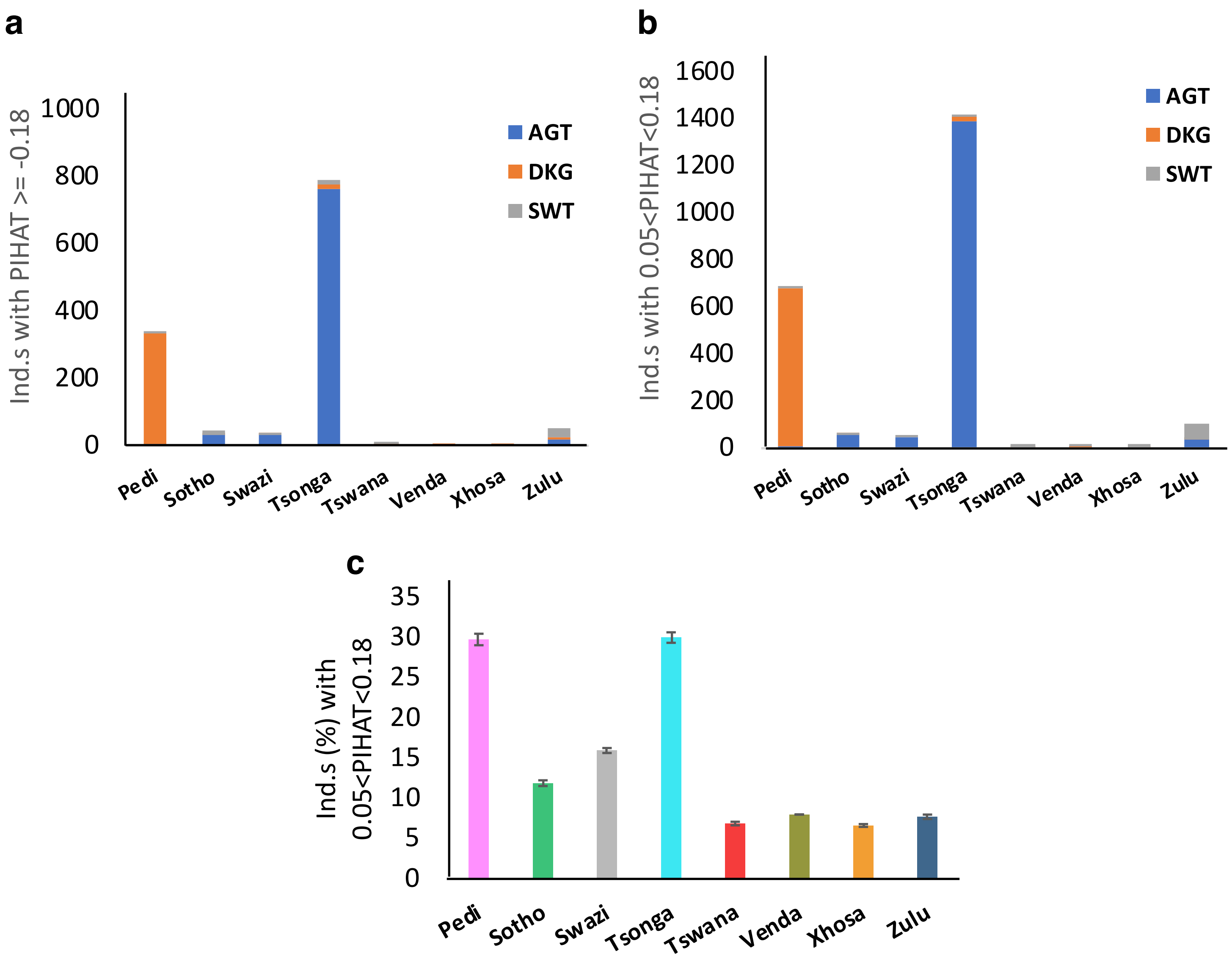


**Supplementary Figure 10. Relatedness levels in South Eastern Bantu-speaking (SEB) groups.** **a**, Relatedness at PIHAT>0.18 in groups stratified by study site **b**, Relatedness at 0.05<PIHAT<0.18 in groups stratified by study site **c**, Cryptic relatedness (CR) estimates (0.05<PIHAT<0.18) based on 100 resampling iterations consisting of up to 100 participants from each group. The plots demonstrate very high levels of CR in Tsonga, sampled predominately from Agincourt (AGT), and Pedi, samples predominately from Dikgale (DKG).

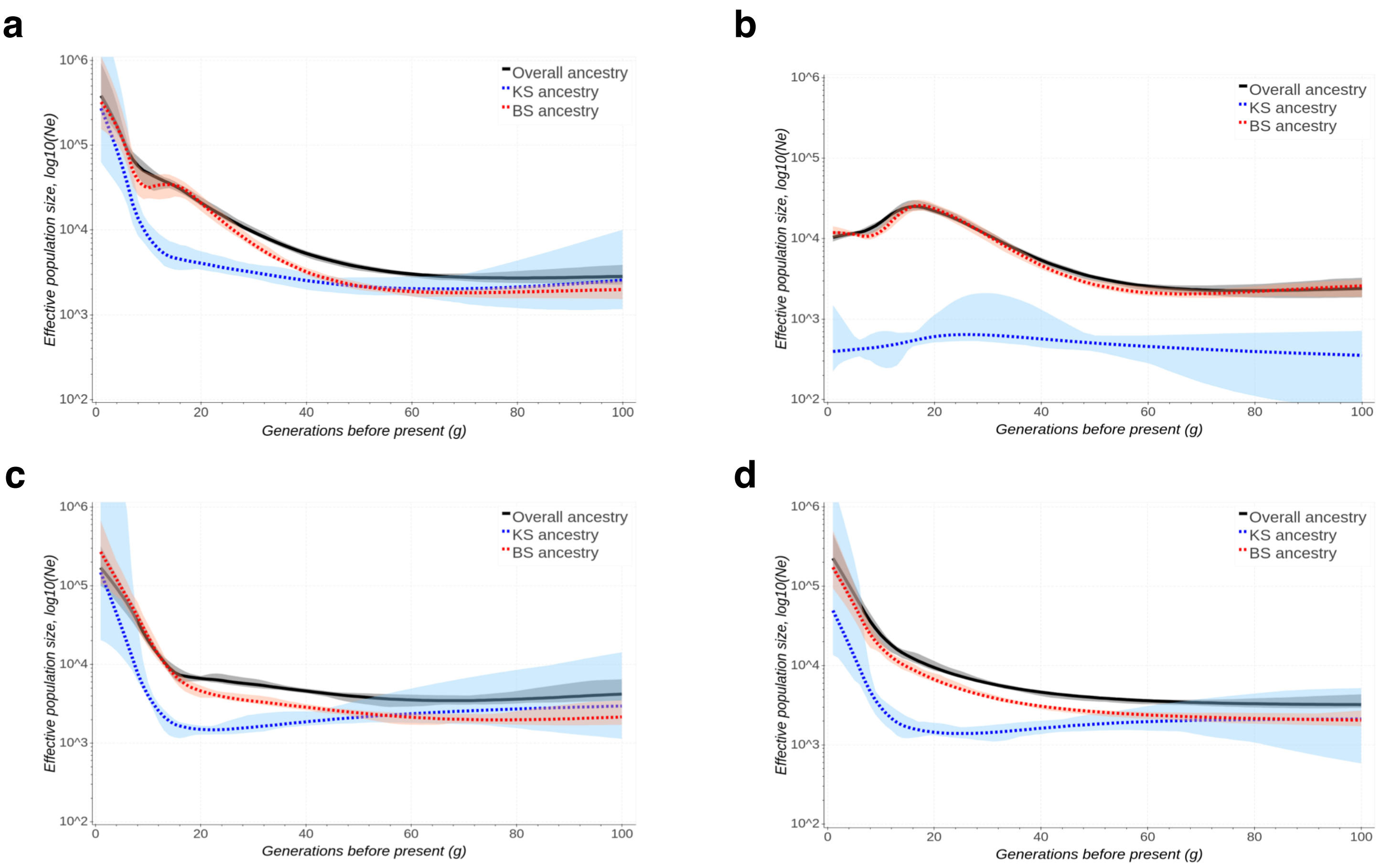


**Supplementary Figure 11 . Khoe-San (KS) and Bantu-speaking (BS) contributions to *Ne* profiles estimated using ASIBD-Ne in four South Eastern Bantu-speaking (SEB) groups: a, Sotho; b, Tsonga; c, Xhosa; and d, Zulu**. The black line shows overall ("true") *Ne* while the red and blue lines shows *Ne* for BS and K-S components, respectively. The shaded areas corresponding to each line demarcate 95% confidence intervals.

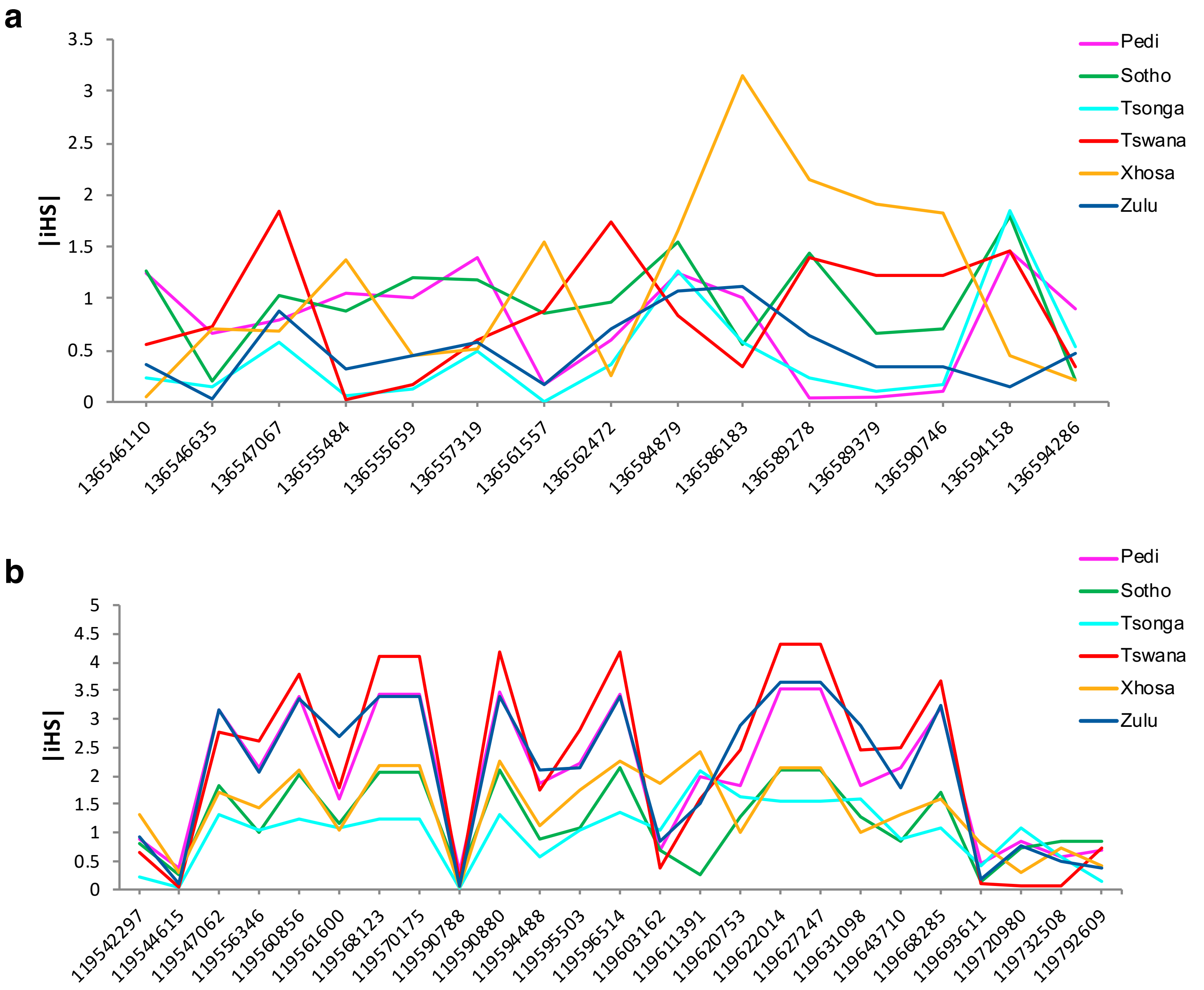


**Supplementary Figure 12. Genomic regions showing high variation in iHS score distribution between South Eastern Bantu-speaking (SEB) groups. a,** *LCT* and **b,** *GSK3B* genes**.** The x-axis in both (a) and (b) shows respective chromosomal coordinates.

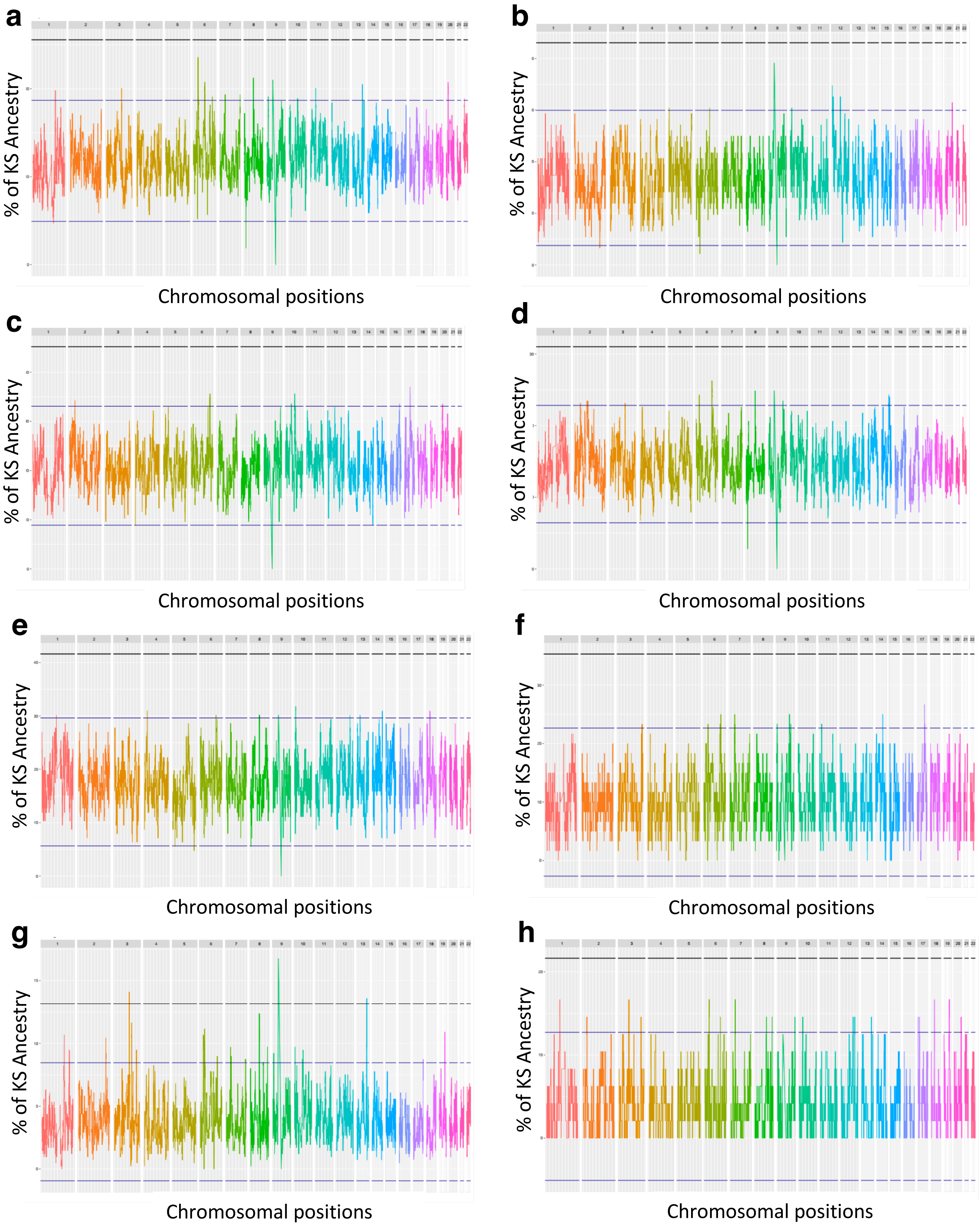


**Supplementary Figure 13. Khoe-San local ancestry distribution across autosomal chromosomes in the South Eastern Bantu-speaking (SEB) group: a, Pedi; b, Sotho; c, Tswana; d, Zulu; e, Xhosa; f, Swazi; g, Tsonga; and h, Venda.** The blue dotted lines represent Mean +/-3SD and the black dotted lines represent Mean +/-6SD.

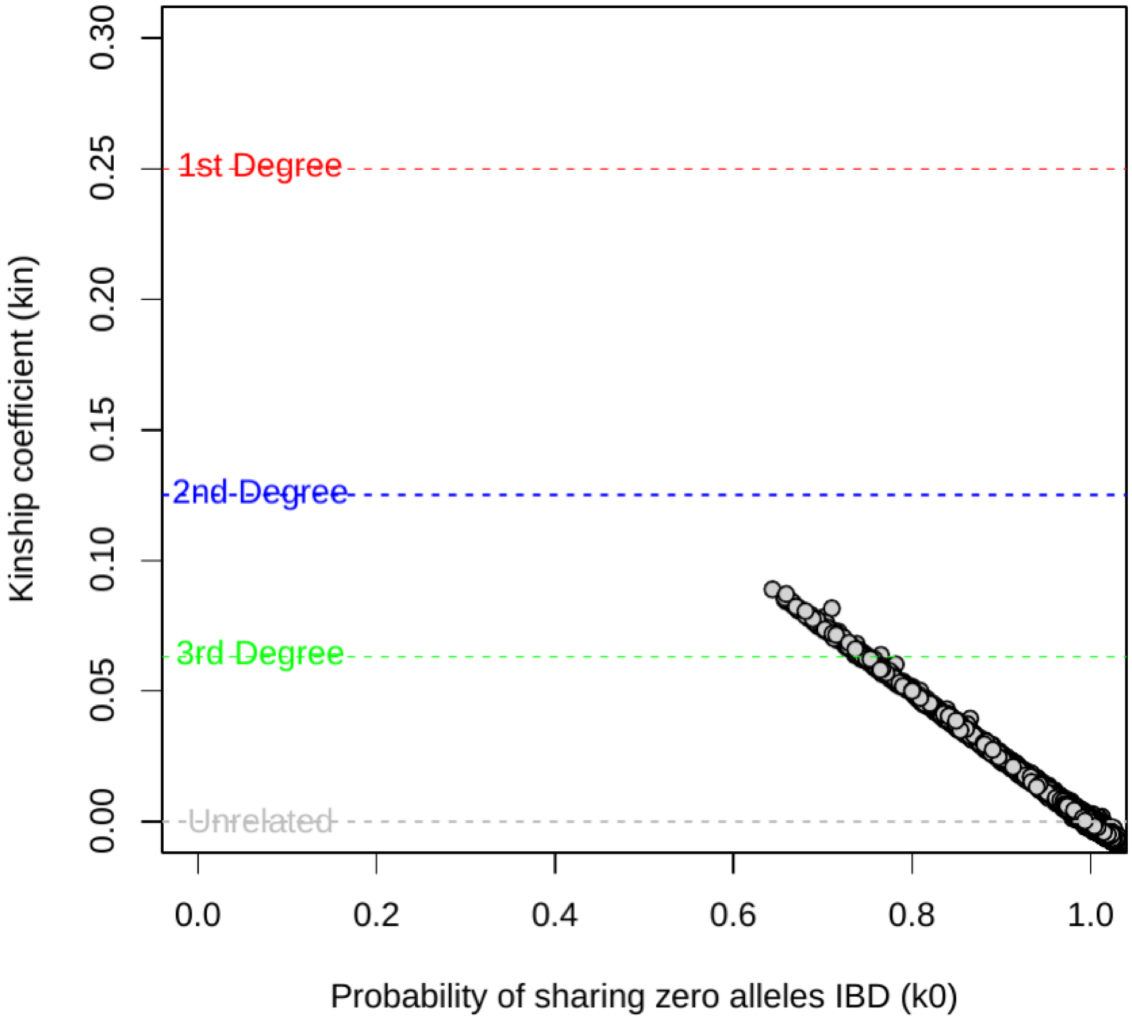


**Supplementary Figure 14. PC-Relate plot depicting measures of pairwise genetic relatedness**. Figure showing kinship coefficient (kin) and probability of sharing zero alleles IBD (k0), obtained using KING and GENESIS. According to the estimated kin and k0 values, low relatedness probabilities between pairwise AWI-Gen samples were found, therefore no first-degree or second-degree relatives were included in subsequent analyses.
