## Supplementary Note for "Genetic-substructure and complex demographic history of South African Bantu speakers"

### SUPPLEMENTARY MATERIAL

#### SUPPLEMENTARY NOTES

##### Note 1. Linguistic phylogeny of the South Eastern Bantu language

In order to compare the genetic relatedness of the populations included in this study with their linguistic affiliation, we include in this paper a new phylogeny of the South-East Bantu (SEB) languages of South Africa, which is part of a larger phylogenetic study of Southern African Bantu languages (Gunnink et al, in prep). The phylogeny is based on lexical data for 100 concepts in 69 Bantu language varieties, 34 of them part of SEB (20 of which are spoken in South Africa) and 35 outgroup languages belonging to different major Bantu branches (ref. [1]). The lexical data were binary recorded in 1,304 partial cognate sets (form-meaning associations). The resulting matrix was analyzed with Bayesian inference methods as implemented in MrBayes (v3.2.7)[2,3] using a restriction-site model[4]. The full majority-rule consensus tree is shown in **Supplementary Fig. 3b**.

Our study confirms that SEB languages are situated within the Eastern branch of Bantu[1]. They descend from a most recent common ancestor that is distinct from other Eastern Bantu languages. Their closest relatives are spoken in the Northeast.

When it comes to internal relationships between SEB languages, our study confirms the well-known split between the groups known as Sotho (including Pedi, Sotho, and Tswana and several smaller varieties not recognized as official) and Nguni (including Zulu, Swazi, Xhosa, Ndebele, and several smaller varieties without official status). Tsonga and its closest relatives in Mozambique form a clade sister to Nguni. Although Tsonga is usually seen as an outgroup, it is according to our phylogeny more closely related to the languages of the Nguni group than to any other South African Bantu language. Together, Tsonga and Nguni constitute a clade that is sister to Sotho. Venda is sister to the clade uniting Sotho with Tsonga and Nguni, which is in line with the language's traditional conception as a relative outsider among South African Bantu languages.

The linguistic phylogeny is mostly in line with findings from genetics and archaeology, but there are also some interesting discrepancies. While the archaeological record suggests that the ancestors of SEB-speaking communities migrated into Southern Africa as already differentiated groups, our linguistic phylogeny provides no evidence for such an early separation. It rather supports a scenario of divergence subsequent to dispersal from a common Southern African homeland, which would be situated in the border region between Zimbabwe, Mozambique and South Africa, based on the principle of highest diversity within SEB. Furthermore, although the identification of genetic Khoe-San (K-S) admixture in certain SEB groups is mirrored in linguistic K-S influence, e.g. in Xhosa and Zulu, other SEB groups show relatively high degrees of K-S admixture yet virtually no linguistic K-S influence, e.g. Tswana[5].

#### Note 2. Dating K-S admixture in SEB groups

To reconstruct the timeframe of admixture events between the major ancestry components in SEB populations, we used three admixture dating methods.

The first method used was fastGLOBETROTTER, the recent implementation of GLOBETROTTER[6]. Briefly, fastGLOBETROTTER tests for evidence of one, two or more pulses of admixture events between two or more ancestral groups, and dates these admixture events to infer the genetic make-up of the studied admixed groups. To do so, we estimated the amount of an individual's genome that is shared with each other individual in the dataset using the chromosome painting approach implemented in ChromoPainter (v4)[7]. This approach applies the model initially introduced by (ref. [8]) to "paint" haplotypically phased chromosomes of a given recipient individual with the haplotypes of all other individuals from other populations included in the dataset. For this analysis, we selected African populations from different African regions and one European population (CEU)[9–11]. Those populations were used as both surrogate and donor populations, and SEB populations as target/recipient populations after randomly down-sampling each SEB population to 30 individuals (except for Venda, in which 24 EC samples were analysed). To estimate program's parameters such as effective population size (*Ne*) and mutation rate (θ), we used ChromoPainter with 10 Expectation-Maximisation (E-M) steps repeating this separately for four chromosomes (1, 6, 12, and 18) and weight-averaging the *Ne* and θ from the final E-M step across the four chromosomes. The estimated values were *Ne*=522.81 and θ=0.00126611, which were used as parameters for "painting" all chromosomes. After chromosomal painting, we used fastGLOBETROTTER to estimate admixture dates for each SEB population following recommendations from (ref. [6]). Confidence intervals (95% CI) of estimates of dates and ancestry proportions were based on 50 bootstrap replicates of the fastGLOBETROTTER procedure.

Second, we used MALDER (v1.0)[12] to test whether a SEB group is admixed between two parental sources (K-S and Bantu-speaker (BS) populations) and estimate the time since admixture based on linkage disequilibrium (LD) decay with distance. All possible triplets of populations in the dataset were tested. To ensure that the varying degree of ancestral components within SEB groups and the difference in sample sizes does not affect the admixture dating, we initially randomly selected a maximum of 100 samples per SEB group in triplicate and ran the analysis. We tested for specific admixture events between each group and K-S hunter-gatherer groups presented in (ref. [11]). The minimum genetic distance to start curve-fitting was set to 0.005 cM to account for short range LD between African populations. Significant results were assessed based on the amplitude of the fitted LD curves and the corresponding z-scores. Concordant results between the 3 analyses were reported. To further test the robustness of the admixture dating, the analysis was repeated with the ethno-linguistically concordant (EC) participants only and yielded similar results.

Third, we used MOSAIC v1.3.7 [13], which exploits admixture LD information to decompose the haploid genome into putative ancestry segments. For each SEB group, we modelled two- and three-way admixture models from unknown ancestral source groups, where the target population is a mosaic of segments from the donor population(s) using a two-layer Hidden Markov Model (HMM) algorithm that allowed for linkage along the haploid genome [13]. This approach can be viewed as a combination of HapMix [14] and GLOBETROTTER[6]. Based on the coancestry plots across individuals in each SEB group and averaged in each group, the best-fitting model was two-way admixture models (with the lowest expected r-squared in all the groups).

To convert the estimated admixture dates from generations to years (on Common Era, CE), we used the formula y= 1950-29*(g+1), where y is the year of admixture, g the estimated number of generations, and taking 29 years as the generation time[15].

The tested admixture models using fastGLOBETROTTER's best-guess conclusion was one date of admixture event in all SEB groups, which is consistent with the ancestry decay curves estimated using MOSAIC on the basis of two-way admixture events in SEB groups. The estimated admixture dates correlated between these two methods (**Supplementary Figure 7b**), except for the Venda group that has more variation in the admixture patterns among the individuals of this group (see SD in **Supplementary Table 2**). In both methods, the estimated source population for the Bantu-related ancestry was Baganda from Uganda[9], and for the K-S-related ancestry were Southern Khoe-San groups, Karretjie and Khomani[11] (**Supplementary Table 2**). Despite similarities in admixture dates, the best K-S proxy populations detected by MALDER were different from the K-S proxy detected by the other two methods (**Supplementary Table 2**). In addition, estimated admixture proportions (BS range: 72-91% and KS range: 9-28%) agree with our previous results using admixture inference methods (**Supplementary Table 2**).

#### Note 3. Sex-specific admixture patterns

Several recent studies based on surveys of mitochondrial DNA (mtDNA) and Y-chromosome (Y-chr) haplogroups in Southern African populations have demonstrated a clear sex-biased gene flow between the K-S and BS[16–18]. Among the five Y-haplogroups found to be common among the SEB of this study, three are associated with Bantu-speakers (E1b1/E-P2, E2b/E-M52, and B2a1/B-M109) and two are associated with K-S populations (B2b/B-P6 and A1b1b2a), which are only 5.1% of the samples (**Supplementary Table 4, Fig. 3a**). Quality of assignment was measured using the F1 score —all assignments of E haplogroups were done with F1 score >0.89, and assignments of B haplogroups were done with F1 score >0.77. The assignment of our samples to A1b was with F1=1, though finer-scale resolution to A1b1b2a was only done with F1 in [0.60, 6.69]. The classification of the relatively few individuals with Y-haplogroups usually not associated with Africans included haplogroups assigned with F1>0.9 except for about a dozen individuals classified in J2a1a with F1<0.6.

In contrast, among the mtDNA-haplogroups detected in our dataset, the proportion of the two K-S associated mtDNA-haplogroups (L0d and L0k) is about 20.5 %, confirming K-S biased maternal gene flow (**Supplementary Table 5, Fig. 3a**). MtDNA classification was more complicated than for Y-haplogroups due to technical limitations of the H3A custom array. Nonetheless, this array allowed high resolution and accurate calling of L0 haplogroups associated with K-S ancestry/speakers (such as L0d and L0k), and could distinguish between three sub-haplogroups of L0d (L0d1, L0d2, and L0d3). However, the base of the array was from existing Illumina bead pools which has good coverage of non-African haplogroups (viz, M and N and below) and some coverage of African haplogroups. As part of the design process, additional probes were added (Botha et al. in prep). However, the underlying array technology probes for SNPs that are within 100 bp of each other may interfere with each other (and more so as they get closer to each other). As the mitochondrial genome is too short (over 16K SNPs) and there were over 200 SNPs genotyped, the array has limitations for the coverage of other African mtDNA-haplogroups. Besides, the classifications that were made were done with reasonable quality scores (except for L2a1 with a score of 0.63), but in some cases it was at a very coarse resolution. For example, 16% of the samples were classified as L0a'b'g but could not be classified more deeply and about 5% were classified as L1'2'3'4'5'6 but could not be classified more deeply. Additional SNPs covering L0a and L0g seem to be the most pressing, and with extra coverage of L3, L4, L6 and especially L2 being desirable.

The comparison of autosomal and X-chromosome contributions in various SEB groups reiterated the overall trend of female-driven gene flow from K-S (**Fig. 3b**). However, as seen with uniparental haplogroup comparisons, the degree of this sex-bias was found to vary widely between the SEB groups, with groups such as Tsonga and Zulu showing a weaker sex-bias in comparison to Xhosa and Sotho (**Fig. 3b**). Sotho shows significantly higher sex-bias in comparison to all the other SEB groups **(Supplementary Table 6).** Moreover, while some groups with higher overall K-S ancestry (Tswana) demonstrate stronger sex-bias in admixture compared to groups with lower K-S ancestry (Pedi), the observed variations in the level of sex-biased admixture are not driven by the differences in K-S ancestry. For example, Zulu in spite of having much higher K-S ancestry in comparison to the Tsonga show comparable sex-biased admixture (*P*-value=0.3). Although our results from both uniparental markers and admixture difference ratio overall support the existing hypothesis of sex-biased admixture between Bantu-speaking males and autochthonous K-S females, the extent of this sex-biased admixture might have varied among SEB groups possibly due to various demographic and cultural factors.

#### Note 4. Levels of relatedness

High levels of relatedness among individuals could potentially influence PCA, admixture profiles and other population-based estimates and need to be accounted for in genome-wide association studies. The assessment of background relatedness in a dataset is, therefore, important for ensuring the robustness of various genetic inferences. Identity-by-descent (IBD) between pairs of individuals from each study site was estimated using PLINK (v1.9)[19]. Pairs of individuals with PIHAT >0.18 were considered to be highly related (equivalent to third degree and closer relationships), while individuals with PIHAT values between 0.05 and 0.18 were considered to show cryptic relatedness (between third degree and fifth degree relatives).

In the AWI-Gen dataset, we estimated that about ~36% of Tsonga participants (predominantly from the Agincourt (AGT)) and ~25% of the Pedi participants (predominantly from the Dikgale (DKG) show a very high relatedness (PIHAT >0.18) (**Supplementary Fig. 10a**). We observed a very similar trend for the number of individuals distantly related to each other (0.05<PIHAT<0.18), where Tsonga and Pedi once again have the highest numbers from AGT and DKG, respectively (**Supplementary Fig. 10b**). To investigate if these observations were biased by the unequal sample size in Tsonga and Pedi, we performed a bootstrap approach by resampling up to 100 samples for 100 iterations, and estimated the percentage of samples related within the range of 0.05<PIHAT<0.18 for each SEB. The analysis reiterated our results, and showed the levels of cryptic relatedness to be high in Tsonga and Pedi (~30 ±1.2 SD%), even after accounting for sample size differences (**Supplementary Fig. 10c**). Moreover, this analysis also demonstrates the cryptic-relatedness levels to be relatively high (range: 10-15%) in some of the other groups such as Swazi and Sotho (**Supplementary Fig. 10c**).

#### Note 5. Signatures of positive selection in SEB groups

We identified regions under positive selection in the SEB by estimating integrated haplotype scores (iHS) for each genic SNP in six SEB groups. All the SNPs that were observed to show extreme outliers iHS scores (|iHS|>4; *P*-value<0.003) in SEB populations are listed in **Supplementary Table 10**. **Fig. 4f** provides a comparison of the distribution of iHS scores for some of the genetic variants that are observed as outliers in at least two of the six SEB groups. As expected, the majority of these variants show uniformly high scores, although not always reaching the outlier threshold across all groups. However, for the outlier variants in genes such as *PAH, CAPN2,* and *SYT1,* the iHS were found to vary more widely between the six SEB groups (**Fig. 4f**). Although these variants emerged as outliers in some SEB groups, no evidence for selection was detected even at a relaxed *P*-value threshold of $P<0.05$ in other SEB groups. Moreover, as iHS can only be estimated for SNPs with a minimum MAF of 0.05, the allele frequencies of the outlier SNPs in genes such as *PPARG, RYR3,* and *SLC8A3* were found to be below this threshold in some of the SEB groups (shown by dark blue in the heatmap) (**Fig. 4f**).

To identify the possible functional impact of the signals, we classified the genes containing outlier SNPs according to ontology, pathway annotations and literature. The major functions represented by these genes include lipid metabolism, circadian regulation, response to oxygen levels, and immune related functions (**Fig. 4f).** One of these immune related genes, *LYAR,* has recently been shown to promote replication of multiple viruses such as influenza A virus (IAV), vesicular stomatitis virus (VSV), Japanese encephalitis virus (JEV), as well as to act as a negative regulator of innate immune responses[20].

Among known African selection signals, only *SYT1* (neurodevelopmental) and *FOXP2* (speech and language) were found to harbour an outlier SNP ($|iHS|>4$). However, we detected signatures of selection around other well-known selected regions, such as *LCT* (Lactase persistence), *LARGE* (Lassa fever), *OCA2* (skin pigmentation), and *VAV*3 (high altitude) at a moderate threshold of $|iHS|>3$ (*P*-value <0.05) (**Fig. 4g**). Interestingly, the signals in the *LCT* gene were found to reach moderate iHS ($|iHS|>3$) only in Xhosa. Moreover, the comparison of the difference in iHS values between SEB groups (**Supplementary Fig. 12)** shows the regions of difference to span a long genomic window (*LCT* in Xhosa) and even an entire gene (*GSK3B* in Tswana and Tsonga). A study (ref. [21]) has shown the presence of a variant in the *LCT* gene, that contributes to lactose persistence (LP) in appreciable frequencies in the Xhosa, and associated this with a high level of K-S ancestry.

To identify variants showing high differentiation between SEB groups, we estimated population branch statistics (PBS) for variants between one pair of SEB groups and the Han Chinese population (CHB; ref. [10]) used as an outlier population (**Supplementary Table 11**). Genetic variants showing longer branch lengths (*P*-value <0.001) in Tswana, when compared to Tsonga, were detected within development related genes (*WLS)*, breast cancer associated genes (*BCSA3* and *BCSA4*) and a gene associated with ebola hemorrhagic fever (*NFKBIE*). The variants showing longer branch length in Tsonga when compared to Tswana, were found in key immune related genes (*VWF* and *ITGB2*), solute carrier genes (*SLC14A2, SLC35E3,* and *SLC1A1*), and the *MCHR1* gene, which play an important role in the control of feeding behaviour and energy metabolism (**Supplementary Table 11**).

#### Supplementary Note References


1. Grollemund, R. *et al.* Bantu expansion shows that habitat alters the route and pace of human dispersals. *Proc. Natl. Acad. Sci. U. S. A.* **112**, 13296–13301 (2015).
2. Ronquist, F. & Huelsenbeck, J. P. MrBayes 3: Bayesian phylogenetic inference under mixed models. *Bioinformatics* **19**, 1572–1574 (2003).
3. Huelsenbeck, J. P. & Ronquist, F. MRBAYES: Bayesian inference of phylogenetic trees. *Bioinformatics* **17**, 754–755 (2001).
4. Felsenstein, J. Phylogenies from restriction sites: a maximum-likelihood approach. *Evolution* **46**, 159–173 (1992).
5. Pakendorf, B., Gunnink, H., Sands, B. & Bostoen, K. Prehistoric Bantu-Khoisan language contact: A cross-disciplinary approach. *Language Dynamics and Change* **7**, 1–46 (2017).
6. Hellenthal, G. *et al.* A genetic atlas of human admixture history. *Science* **343**, 747–751 (2014).
7. Lawson, D. J., Hellenthal, G., Myers, S. & Falush, D. Inference of population structure using dense haplotype data. *PLoS Genet.* **8**, e1002453 (2012).
8. Li, N. & Stephens, M. Modeling linkage disequilibrium and identifying recombination hotspots using single-nucleotide polymorphism data. *Genetics* **165**, 2213–2233 (2003).
9. Gurdasani, D. *et al.* The African Genome Variation Project shapes medical genetics in Africa. *Nature* **517**, 327–332 (2015).
10. 1000 Genomes Project Consortium *et al.* A global reference for human genetic variation. *Nature* **526**, 68–74 (2015).
11. Schlebusch, C. M. *et al.* Genomic variation in seven Khoe-San groups reveals adaptation and complex African history. *Science* **338**, 374–379 (2012).
12. Loh, P.-R. *et al.* Inferring admixture histories of human populations using linkage disequilibrium. *Genetics* **193**, 1233–1254 (2013).
13. Salter-Townshend, M. & Myers, S. Fine-Scale Inference of Ancestry Segments Without Prior Knowledge of Admixing Groups. *Genetics* **212**, 869–889 (2019).
14. Price, A. L. *et al.* Sensitive detection of chromosomal segments of distinct ancestry in admixed populations. *PLoS Genet.* **5**, e1000519 (2009).
15. Fenner, J. N. Cross-cultural estimation of the human generation interval for use in genetics-based population divergence studies. *Am. J. Phys. Anthropol.* **128**, 415–423 (2005).
16. Bajić, V. *et al.* Genetic structure and sex-biased gene flow in the history of southern African populations. *Am. J. Phys. Anthropol.* **167**, 656–671 (2018).
17. Schlebusch, C. M. Genetic variation in Khoisan-speaking populations from southern Africa. (University of the Witwatersrand Johannesburg (South Africa, 2010).
18. Choudhury, A. *et al.* Whole-genome sequencing for an enhanced understanding of genetic variation among South Africans. *Nat. Commun.* **8**, 2062 (2017).
19. Chang, C. C. *et al.* Second-generation PLINK: rising to the challenge of larger and richer datasets. *Gigascience* **4**, 7 (2015).
20. Fang, Y. *et al.* New strains of Japanese encephalitis virus circulating in Shanghai, China after a ten-year hiatus in local mosquito surveillance. *Parasit. Vectors* **12**, 22 (2019).
21. Ranciaro, A. *et al.* Genetic origins of lactase persistence and the spread of pastoralism in Africa. *Am. J. Hum. Genet.* **94**, 496–510 (2014).
22. Semo, A. *et al.* Along the Indian Ocean Coast: Genomic Variation in Mozambique Provides New Insights into the Bantu Expansion. *Mol. Biol. Evol.* **37**, 406–416 (2020).
